# Systematic discovery of protein kinase-like domains reveals diverse evolutionary strategies in the human oral microbiome

**DOI:** 10.64898/2026.09.18.752743

**Authors:** Marianna Krysińska, Weronika Śliczniak, Zuzanna Jakubowska, Marcin Gradowski

**Author notes:** Correspondence: Marcin Gradowski.

## Abstract

The protein kinase-like (PKL) superfamily regulates metabolism, biofilm formation, host interactions, and antimicrobial resistance in bacteria, although much of its diversity remains unknown. We used a structure-based pipeline that combined protein structure prediction with structural similarity searches to search 5.1 million protein sequences in the Human Oral Microbiome Database (HOMD). Together with 30 families that had been previously characterized, we identified 20 PKL families new to this genome collection: 15 entirely novel and 5 previously reported as preliminary findings. Our taxonomic analysis revealed that the families were distributed either across multiple bacterial phyla or restricted to a single species; hits spanning domains suggest either ancient origins or horizontal transfer. Three families were examined in detail and show different evolutionary pathways: SEAE1 from *Segetibacter aerophilus*, which is a putative lipid kinase; POGI1 from *Porphyromonas gingivalis*, a putative ethanolamine kinase with the PKL fold that is limited to pathogens; and SrfA-N from *Haemophilus parainfluenzae*, a non-catalytic scaffold that has been convergently co-opted as a tripartite toxin platform. Taken together, these examples demonstrate enzymatic specialization, adaptation specific to pathogens and structural co-option. The families identified are candidates for further mechanistic study.

## Introduction

The human oral microbiome (HOM) is a complex and dynamic community of microorganisms that lives in the oral cavity; it is one of the most diverse environments in the human body and plays a vital role in both oral health and disease [1].

HOM consists of bacteria, fungi, viruses, and protozoa, with bacterial species being the most numerous and the subjects of the most of research. About 1,000 species or phylotypes have been identified, these belonging to the phyla Actinobacteria, Bacteroidetes, Chlamydia, Fusobacteria, Firmicutes, Proteobacteria, Spirochaetes, and Tenericutes. Moreover, approximately half of the bacterial species found in the mouth are difficult to cultivate [1,2].

The composition of HOM is significantly affected by diet, oral hygiene habits, smoking, and other lifestyle factors. For instance, diets high in sugar encourage the growth of bacteria that cause tooth decay. When there is an imbalance in the HOM, a condition known as dysbiosis, this is connected with oral diseases. Bacteria such as *Streptococcus mutans* are linked to tooth decay [1,3], and other bacteria, such as *Porphyromonas gingivalis*, are involved in periodontal disease [4]. It is increasingly important to understand individual differences in HOM to provide personalized dental care and predict susceptibility to oral diseases. In addition to periodontitis, dysbiosis has been linked to systemic diseases. Chronic inflammation can occur when dysbiosis results in infection of the periodontal pocket by pathogens. The inflammation thus causes cytokines to be released, and these spread through the bloodstream, possibly leading to systemic inflammation and thus raising the risk of various conditions such as stroke, colon cancer, obesity, diabetes, atherosclerosis [5], and Alzheimer’s disease [6].

Understanding of the HOM has been greatly improved by advances in genomic and metagenomic analyses, with current studies examining interactions between oral microorganisms and the roles these microbes play in health and disease [7].

Protein kinases are regulatory enzymes that affect cellular process through protein phosphorylation. The so-called protein kinase-like (PKL) superfamily includes catalytically active kinases, catalytically inactive pseudokinases that have regulatory functions, and kinase-fold proteins that show a variety of enzymatic activities [8–11]. In oral microorganisms, these enzymes control metabolic pathways, are involved in the response to environmental stress, help in the formation of biofilms, and allow adaptation to changing conditions [12–14]

In this study, the HOMD, the most comprehensive source of genomic and metagenomic data on microorganisms inhabiting the human oral cavity [15], was used in a sequence and structural bioinformatics study to identify PKL proteins. A survey of HOMD against established PKL family profiles [8] first confirmed the presence of 30 previously characterized PKL families. This mainly comprises eukaryotic-type protein kinases [16,17]; choline kinases which initiate the addition of phosphorylcholine to teichoic acids [18–20]; and the lipopolysaccharide (LPS) kinases which belong to the Kdo family and phosphorylate the inner core of LPS in Gram-negative bacteria [21]; aminoglycoside and macrolide phosphotransferases that confer antibiotic resistance [22]; and HipA-family toxins associated with the formation of persisters [23,24] (Figure 1; Suppl. Data 1 Histogram). Building on the previous approach [9], 20 PKL families were identified in the HOMD. Three of these (LACHF, LACHP, CHLO1) were preliminarily reported as part of the KINtaro database development [8]. For two additional families (TREV1, TRSO1), representative proteins were analyzed in an earlier MSc thesis [25]. All five are defined and characterized at the family level here for the first time; the remaining 15 represent novel discoveries.

**Figure 1.**
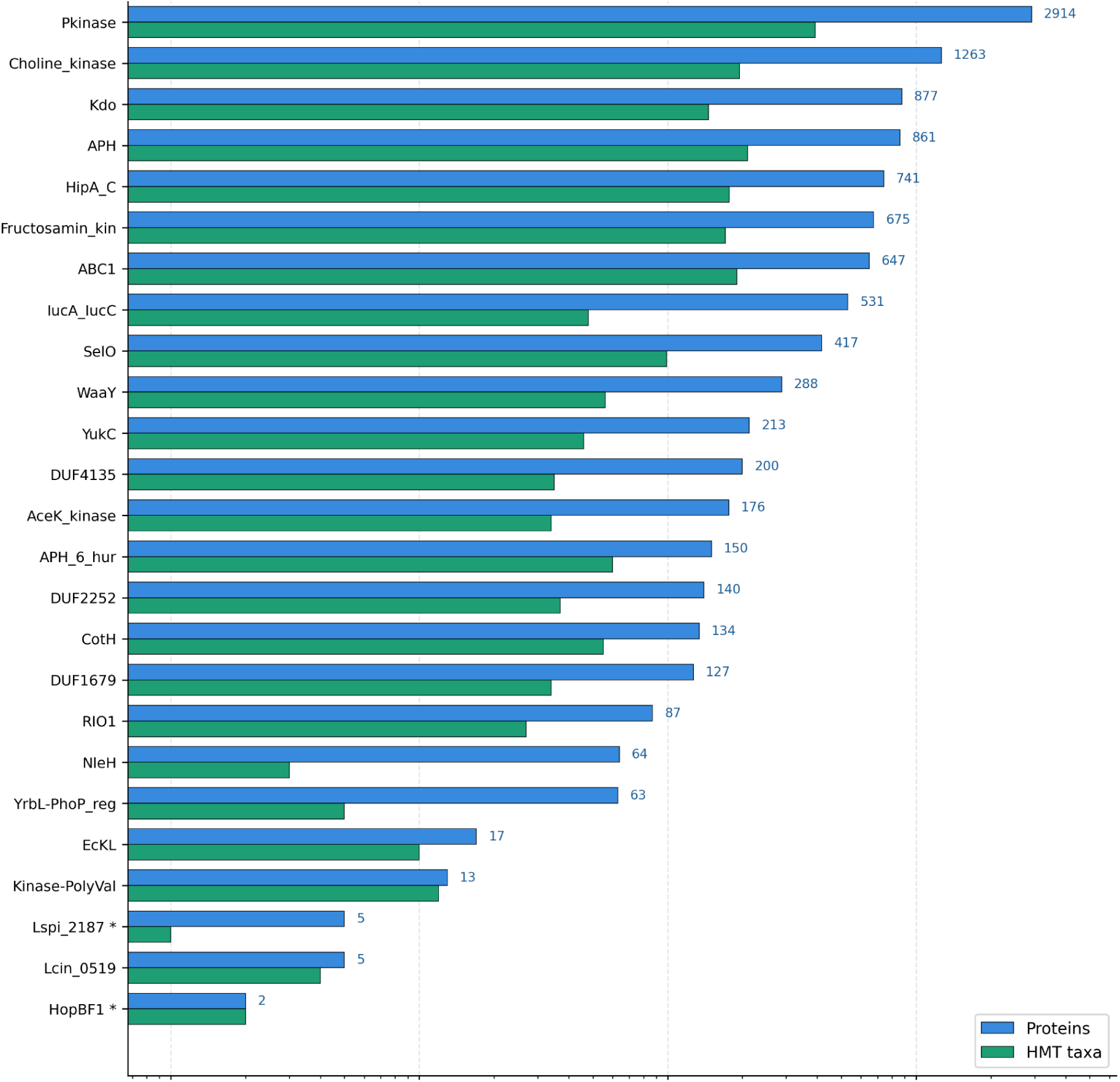
Previously characterized PKL families were detected in the human oral microbiome. Proteins (blue) and HMT taxa (green) are assigned to each PKL family in HOMD v9.15a, shown on a logarithmic x-axis. Of the 30 families reported (see Methods), 25 with more than one protein are shown and are ordered by protein count. Five singletons are left out for clarity. Asterisks indicate families listed as provisional. Another provisional family, Act-Frag_cataly, is a singleton and is not shown (Figure 1; Suppl. Data 1 Histogram).

## Results and discussion

In order to identify new PKL proteins systematically, a structure-based discovery pipeline was used to analyze 5.1 million protein sequences from the HOMD (v9.15a) [14,27]. The sequences were clustered, fragmented, and known domains were removed before protein modeling was performed with ESMFold. Structural comparisons with the established PKL families were made using TM-align, HHpred, FATCAT and DALI [28–31] As a result, 20 PKL families new to HOMD were identified (Suppl. Table S1; Suppl. Table S2, thus increasing the known variety of kinase-like proteins found in bacteria of the oral microbiome. Three of these families had been reported previously during the development of the KINtaro database; the other 17 are discoveries that are specific to this study [8].

### Structural versus sequence-based network analysis of PKL families

Graph analysis (Figure 5) enables assessment of sequence and structure similarities between 17 newly identified PKL HOM families and 71 established PKL families from all domains of life. This graph highlights potential distant relationships between families, which are critical for understanding their evolutionary trajectories and functional roles. The classic CLANS graph depicts quasi-distances between sequences derived from “all-to-all” sequence comparisons performed with BLAST [33]. In addition, we used the NISARA script, which, analogously, determines quasi-distances between structures based on TM-align comparisons (transformed TM-scores) (Suppl. Table S3; Suppl. Data 3) [34].

We have deliberately used this method because classical phylogenetic methods depend on a reliable multiple sequence alignment, which is difficult to obtain for highly diverse and distantly related families due to insertions and deletions specific to each family. Similar difficulties arise with multiple structural alignment faces, analogous challenges at this level of divergence, as no single reference frame can accommodate all superfamily members simultaneously. The approach used in NISARA, which is pairwise structure comparison, overcomes this limitation by considering each pair separately.

Analysis reveals that the NISARA network has a single PKL core, whereas the sequential CLANS network breaks down into several families. This shows that there is high sequence variability coupled with a conserved structural backbone.

In the sequence-based graph, most novel families are at the periphery. In contrast, in the structural CLANS, many novel families are near the center of the PKL structure cluster. Only the CODI1 and LACHP family structures are atypical.

We highlight several families that illustrate the complementarity of both approaches: only their combination allows us to grasp non-obvious similarities.

The TREV1 and TRSO1 pseudokinase families are closely grouped in the structural CLANS, but form separate clusters in the sequence-based analysis. This suggests a shared PKL fold with significant motif divergence.

Although the CACU1 pseudokinase family is near the Pox-ser/thr cluster in the structural diagram (with no connections in the sequential diagram), it shows the strongest direct structural similarity to NleH, rather than to Pox-ser-thr-kin.

In CLANS, DUF2252 is connected to the PKLs but is isolated in NISARA, probably due to a limitation of TM-align. Since TM-align maintains the order of residues [28], a permutation of the N and C lobes can lead to a reduction in the TM-scores, whereas BLAST still identifies local conserved kinase motifs [27,32]. Thus, this discrepancy reflects method bias rather than true fold divergence.

Further examples highlight the complementarity of both approaches. NIRAN is positioned close to the PKL core in the structural network, yet appears at the periphery in the sequence-based graph. Although NIRAN is situated near the center of the structural network, it is found at the edge in the sequence-based graph, while Pox-E2-like and IucA-IucC retain strong structural links to the core, which are mostly missing from the sequence representation. These cases support the general trend that structural remnants of the PKL fold continue to exist even in the face of considerable sequence divergence, and it is only by combining both network and graph types that the full range of relationships among the members of the superfamily can be captured.

### Three evolutionary trajectories of the PKL fold

The 20 identified PKL families exhibit substantial variation in taxonomic breadth, domain architecture, and conservation of catalytic motifs (Figure 3; Suppl. Table S1; Suppl. Logo; Suppl. Data 2). Comprehensive characterization of all families is beyond the scope of this study. Instead, three families were selected for detailed bioinformatic analysis to represent the observed spectrum of PKL diversification, rather than the most abundant or conserved members. SEAE1 demonstrates structural similarity (Suppl. Table 2) to PI3Ks and is broadly distributed across bacteria. The PKL fold is observed in POGI1, which is restricted to two strains of a single oral pathogen and has both a complete active site and an ethanolamine-kinase-like environment, indicating a pathogen-specific adaptation. SrfA-N, which is found in a wide range of Gram-negative bacteria, is an example of the PKL scaffold being used for a non-enzymatic function. Together, these three examples show different evolutionary paths: enzymatic specialization, pathogen-specific adaptation, and structural co-option, all of which can occur within a single microbial habitat.

**Figure 2.**
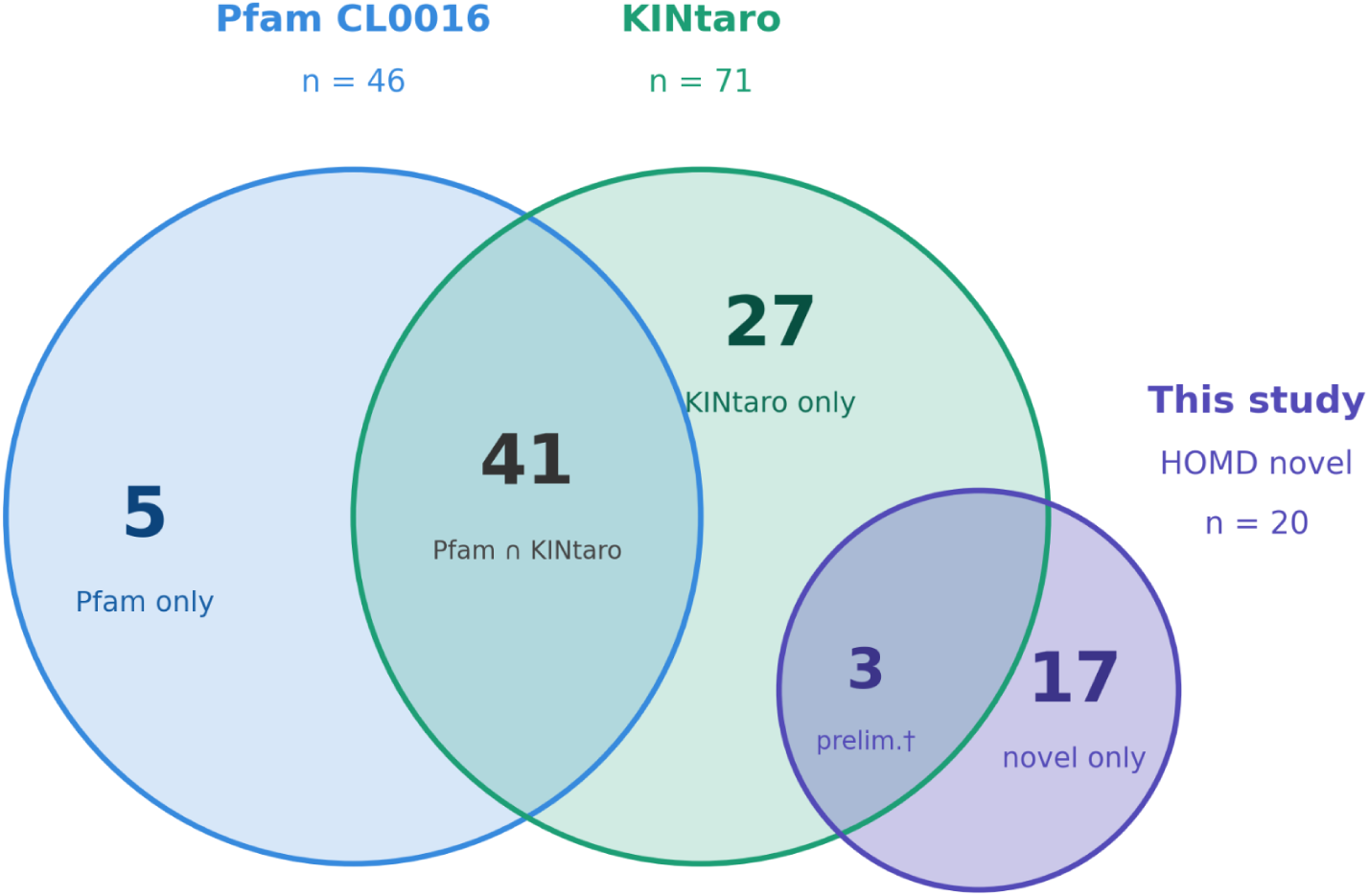
Comparison of PKL family representation across Pfam, KINtaro, and this study. Venn diagram (Suppl. Data 1 venn) of protein kinase-like families cataloged in Pfam clan CL0016 (n=46), KINtaro (n=71), and identified in this study from the human oral microbiome (n=n=20; 17 outside KINtaro). Numbers indicate counts per region. † LACHF, LACHP, and CHLO1 were preliminarily reported in KINtaro [8]. LACHF and LACHP were initially described in [26]. Representative proteins of TREV1 and TRSO1 were analyzed in an MSc thesis [25].

**Figure 3.**
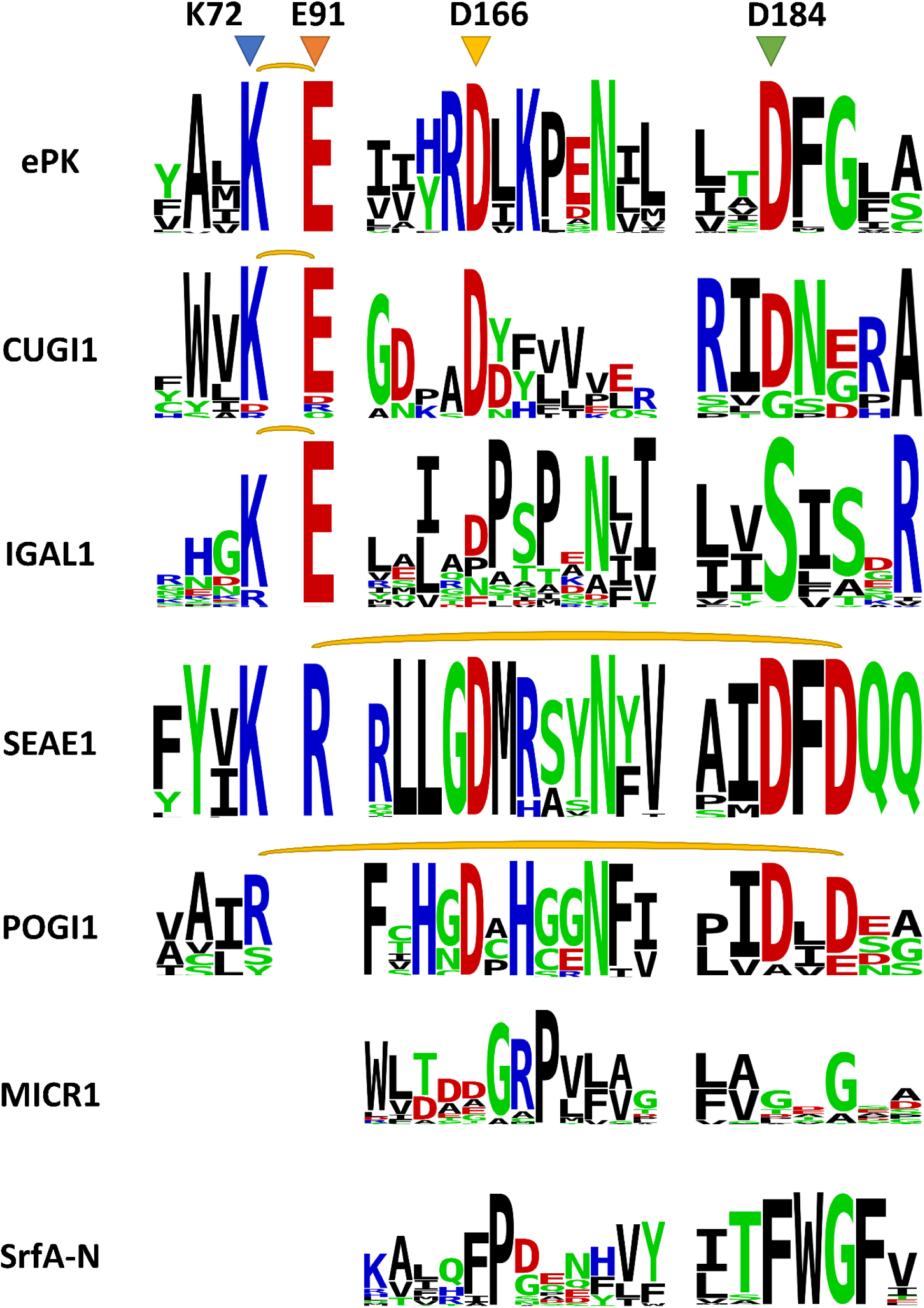
Selected HOMD kinase-like families with conserved active-site motifs. Sequence logos of active site motifs for selected families - logo generated by LogoSS. Also, the “classic” kinases (ePK - generated by sequence of 1ATP - PKA [27]) are shown. Residue numbering (top row) according to standard protein kinase A (PKA) nomenclature. For some families, it was not possible to identify the residue corresponding to ion pairs. Yellow brackets indicate (predicted) salt bridge [27]. MICR1 and SrfA-N have degraded motifs.

### A PAP2–PIK-like phospholipid homeostasis module in *Segetibacter aerophilus*

A novel PKL-family putative lipid kinase, WP_147203046.1 (“SEAE1”), was identified in *Segetibacter aerophilus* strain NBRC 106135. The genome of this strain is represented in HOMD, although the strain itself was originally isolated from air rather than from an oral source [37]. *S. aerophilus* is a Gram-negative, strictly aerobic rod belonging to the family *Chitinophagaceae*. Airborne or dust-derived taxa such as *Chitinophaga*, *Sphingomonas*, or *Segetibacter* are frequently detected in saliva, dental plaque, and oral biofilm datasets; however, they typically represent environmental background rather than indigenous members of the oral microbiome.

The BLAST homology analysis found 7,867 sequences similar to SEAE1, most of which were located in the Bacteroidota (39.1%) and Verrucomicrobiota (8.7%), with a small number in the Archaea (0.4%). The fact that these sequences are so sparsely distributed across different domains is compatible with either an origin that predates the divergence of the bacteria and archaea or with horizontal gene transfer; to tell which of these two scenarios is correct would need a phylogenetic analysis of representative sequences [38]. The distribution pattern suggests that SEAE1 arose in free-living bacterial lineages and was subsequently retained in taxa associated with host surfaces [39].

Sequence-based similarity to PF00454.33 (PI3_PI4_kinase) is weak (HHpred E-value = 250, Suppl. Table 1D) [29], but structural comparisons always identify lipid kinases as the closest previously characterized relatives (Suppl. Table 2). It is noteworthy that members of the Bacteroidota phylum - such as *Segetibacter*, *Flavobacterium*, and *Sphingobacterium* - synthesize inositol-containing phospholipids even though they do not have classical eukaryotic membranes [40], providing a plausible substrate context for a PKL-fold lipid kinase in this lineage.

AlphaFold3 structure models of SEAE1 were built with ATP (see Suppl. Data 4A). The catalytic core of SEAE1 retains the typical basic and acidic residues required for phosphotransfer: K140 (K72 in PKA), D216 (D166), and D241 (D184). K140 coordinates the α-phosphate and R146 the β-phosphate of ATP. D216 is positioned to act as a catalytic base. Notably, the AF3 model does not show a canonical β3-K72-αC-E91 ion pair; instead, R146 (on the αC-equivalent helix) forms a salt bridge with D243 (2.7 Å), which we interpret as a structurally equivalent, non-canonical substitute for this regulatory ion pair. Catalytic competence is further supported by canonical coordination of two Mg²⁺ ions: N221 stabilizes the first Mg²⁺ position and D241 the second [27,36].

Secretion system analysis of SEAE1 homologs (Suppl. Table S3A) confirmed intracellular localization, with all bacterial sequences classified as non-secreted. In *S. aerophilus*, only the presence of a type IX system is predicted, associated with periplasmic transport rather than effector secretion [41].

In the *Segetibacter aerophilus* genome (accession GCF_049949245.1), the gene encoding SEAE1 (WP_147203046.1) resides within a conserved gene cluster (Suppl. Table 5A, Suppl. Data 4A - OperonMapper [42]) comprising a PAP2-family phosphatase, a secreted/periplasmic protein of unknown function (WP_147203042.1), the tmRNA-binding protein SmpB, the acetyl-CoA carboxylase β-subunit (AccD), and SEAE1 itself. While the SEAE1 domain is broadly distributed across bacterial phyla (7,867 homologs), this specific gene-neighborhood arrangement is restricted to the *Segetibacter* lineage: it is intact in 4 of 20 currently available *Segetibacter* genomes, whereas the remaining 16 retain only fragmented remnants of the locus, which may partly reflect assembly incompleteness. This suggests that the coupling of SEAE1 with a lipid phosphatase into a putative functional module represents a lineage-specific arrangement rather than an ancestral feature of the domain.

The PAP2-family phosphatase is a membrane hydrolase that removes phosphate groups from phosphatidic acid, phosphorylated lipids, and related metabolites, consistent with a role in membrane remodeling and phospholipid homeostasis [43]. Sequence WP_147203042.1 contains a strong Sec/SPI signal peptide (Suppl. Table S5A), indicating secretion to the periplasm [44]; no lipobox is present, and all HHpred hits show weak support, so this protein is annotated as a secreted or periplasmic protein of unknown function without a reliable domain assignment (Suppl. Table S5A) [29]. SmpB forms a complex with tmRNA and participates in trans-translation, a rescue mechanism that prevents ribosome stalling on damaged mRNA [45,46]. AccD is the β-carboxyltransferase subunit of the acetyl-CoA carboxylase complex, a key enzyme of fatty acid biosynthesis [47,48]. Operon prediction indicates that these genes are not co-transcribed (see Suppl. Data 4A - OperonMapper), which means their arrangement probably reflects the conserved genomic proximity of genes with related functions rather than indicating the presence of a coordinately regulated operon.

Genomic neighborhood analysis using WebFLAGS (Suppl. Data 4A - webFlaGs_SEAE1) shows that other SEAE1 homologs - outside the *Segetibacter*-specific cluster - co-occur with genes encoding helical-backbone receptor/ABC-transporter substrate-binding proteins, consistent with a broader association with transport- or membrane-related functions. In some genomes, class I/II fructose-bisphosphate aldolase genes are also found nearby, suggesting a possible link between carbohydrate metabolism and membrane lipid remodeling across the wider homolog set [40,49–51].

In summary, SEAE1 is a previously undescribed PKL-fold protein with an intact catalytic core, including canonical ATP- and Mg²⁺ coordinating residues and a structurally equivalent substitute for the regulatory ion pair found in active protein kinases [27]. SEAE1 is embedded within a *Segetibacter*-specific gene cluster that also includes a PAP2-family lipid phosphatase. However, inositol lipid biosynthesis has so far been characterized only in host-associated Bacteroidetes lineages via a dedicated gene cluster in *Bacteroides thetaiotaomicron* [40,52]. The presence of a lipid phosphatase in the immediate genomic neighborhood raises the possibility that SEAE1 acts within phospholipid rather than protein metabolism. Whether the environmental Bacteroidota, including Segetibacter, share this metabolic capacity has not been established. The substrate and physiological role of SEAE1 remain undetermined and will require experimental characterization. SEAE1 thus illustrates how the PKL fold can be retained with a fully active-looking catalytic core while being redeployed, potentially, toward a non-protein substrate.

### Novel PKL-fold kinase predicted to drive *P. gingivalis* virulence

In the oral strain *Porphyromonas gingivalis* TDC60, we identified the protein POGI1 (WP_013815849.1) as a remote homolog *of* known protein kinase families (Suppl. Table S1/2). The TDC60 strain exhibits greater pathogenicity than W83 and ATCC 33277 in a mouse abscess model [53]. A clear homolog of POGI1 is only found in strain KCOM 3131, for which essentially no experimental data are available, suggesting species-specific pathogenic adaptation (Suppl. Table S3B). Three distant homologs are present in poorly characterized *Archaea*. *P. gingivalis* is an anaerobic gram-negative bacterium, considered one of the key pathogens in the pathogenesis of chronic periodontitis. Its presence is associated not only with local inflammation, but also with several systemic diseases, including cardiovascular disease, type 2 diabetes, and even Alzheimer’s disease [54].

POGI1 exhibits a PKL-like fold but with a non-canonical arrangement of its active site. Unlike ordinary kinases, it lacks the usual K72-E91 ion pair and instead has R49 in strand beta-3 (see Suppl. Data 2); this residue is 4.5 Å from D386 in the C-lobe. The important catalytic residues are D299 (D166, catalytic loop) and D384 (D184, activation loop), which bind Mg²⁺, as well as the conserved N304 (N171) that coordinates a second metal ion; W44 stabilize the ATP ring. The AlphaFold3 model was created using ATP and Mg²⁺ and yielded an ipTM value of 0.94, indicating a high level of confidence (Table 2). R26 interacts with the α-phosphate, while K242 interacts with the β-phosphate. R266 and H301 are likely involved in bending and positioning the γ-phosphate [27,36,55]. The extended distance between the ion pair may represent a substrate-induced conformational switch mechanism. Salt-bridge formation between these residues can regulate kinase activity by modulating Mg2+ binding [56]. The observed 4.5 Å separation between R49 and D386 suggests that POGI1 was captured in an open or pre-catalytic conformation, similar to the conformational states observed in protein kinase A during catalytic turnover [57]. Although the HHpred hit to cd05157: ETNK_euk (Eukaryotic Ethanolamine kinase) is weak (E-value = 0.66), Suppl. Table 1D) [29]. Profile HMM-based annotation (OperonMapper) independently assigned POGI1 to COG0510 (predicted choline kinase involved in LPS biogenesis), further supporting a putative ethanolamine/choline kinase function. However, the per-sequence E-value for this assignment is not directly reported by the tool [42]. Structure-based comparisons clearly support a PKL/APH-like fold (Suppl. Table S1D/2) [29–31]. The ipTM for the ATP/Mg²⁺-bound model is ∼0.94 [36,58]. The substrate pocket is small and polar, which is compatible with the binding of small polar molecules. Since there is a weak hit with cd05157 (ETNK_euk), POGI1 may act as an ethanolamine kinase and thus initiate the CDP-ethanolamine (Kennedy) pathway when it is part of such operons [29,59,60].

AlphaFold3 was employed to model both the neutral and protonated forms of ethanolamine (Suppl Data. 4B Table 1). The neutral form (Figure 7) exhibited slightly greater positional coherence, as indicated by a lower min PAE, compared to the protonated form, which displayed an increased min PAE. Both forms are compatible with a compact, polar pocket, although there is a preference for the neutral ligand [61]. In conjunction with the non-canonical active-site features replacing K72, the absence of a canonical E91 partner, and the proximity of C-lobe D386, these findings suggest that the protein may function as a pseudokinase, serving as an ATP/ligand-binding regulator rather than a catalytic ethanolamine kinase [10,27].

**Figure 4.**
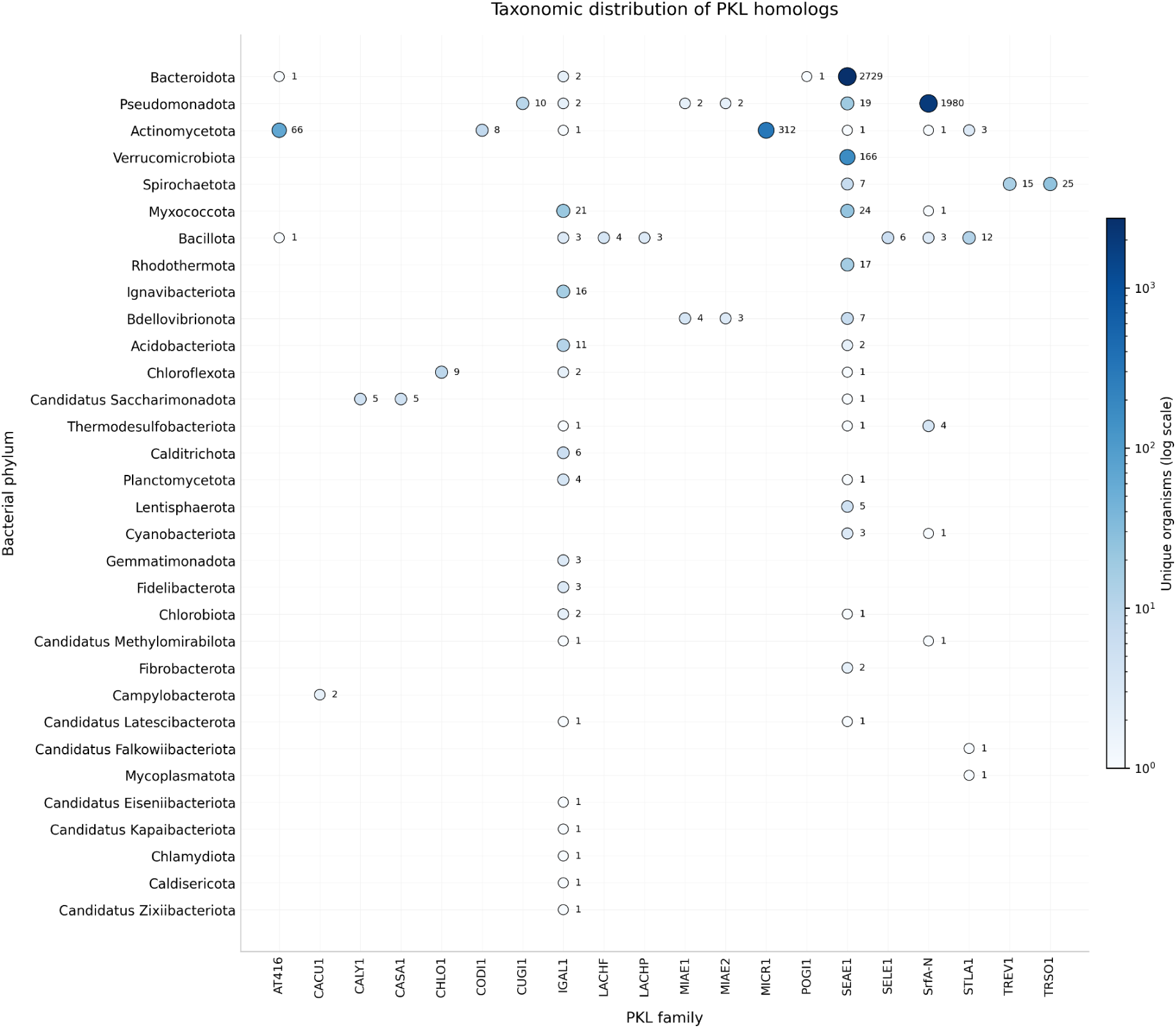
Taxonomic distribution of HOMD PKL families. A dot plot showing the number of unique organisms per PKL family (on the x-axis) for the various bacterial phyla (on the y-axis); the intensity of the colour corresponds to the number of homologs on a log₁₀ scale (as indicated by the colour bar), with numerical labels giving the exact counts; empty cells show that no homologs were detected; archaeal hits are excluded (Suppl. Data 1 dotplot) [32].

**Figure 5.**
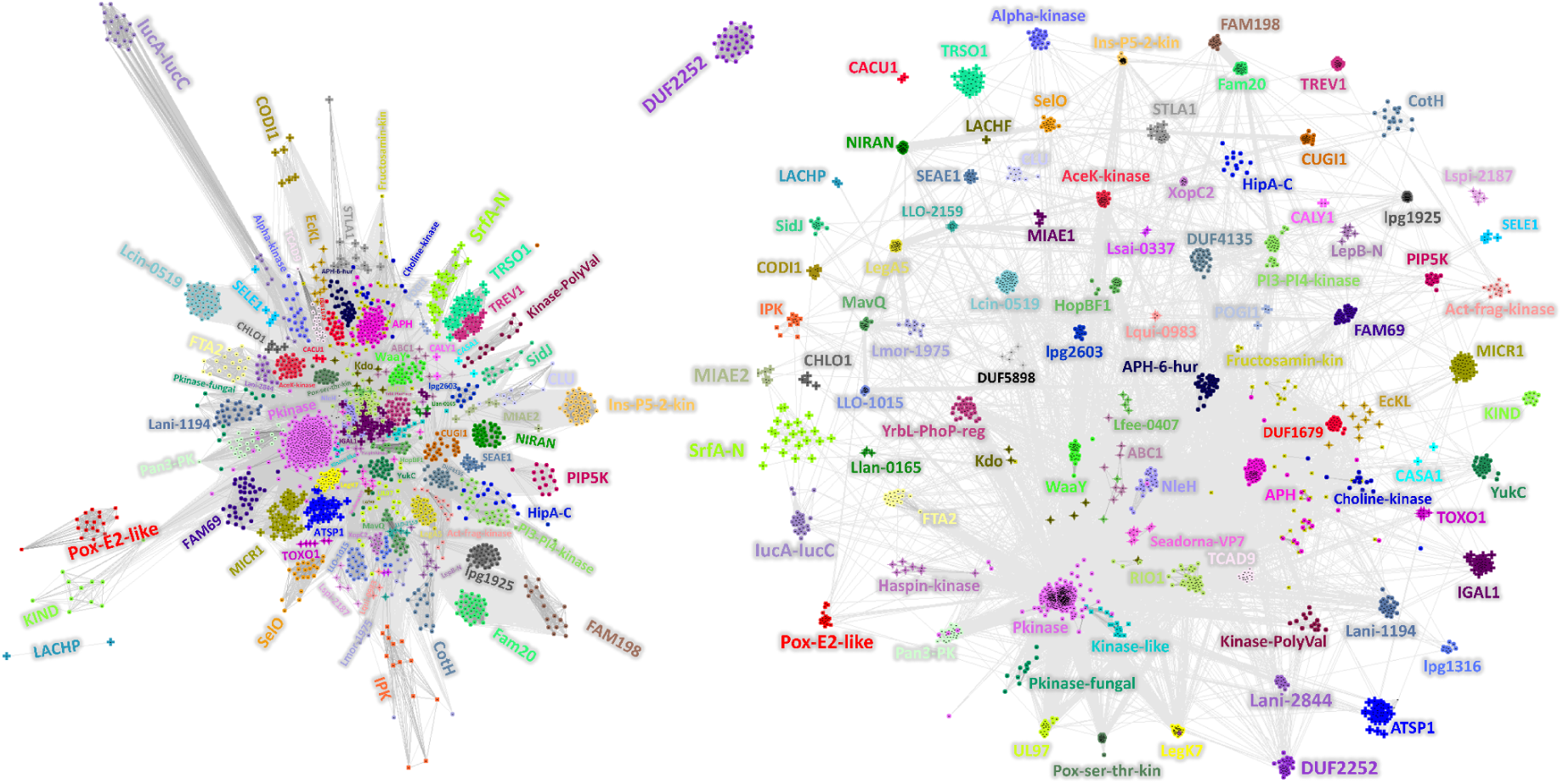
NISARA clustering of structures (left) and CLANS of sequences (right) for the PKL superfamily [33,34]. Plus signs “+” indicate newly described families. Dots “●” and stars “✦” indicate known families. Lines are drawn for pairs that meet the threshold. Proximity in space results from the aggregation of multiple edges (transitive effect) and therefore does not automatically imply homology. In NISARA structure clustering, the threshold cutoff (th-cu) = 0.4, in CLANS of sequences, BLAST HSPs with E-values up = 1.

**Figure 6.**
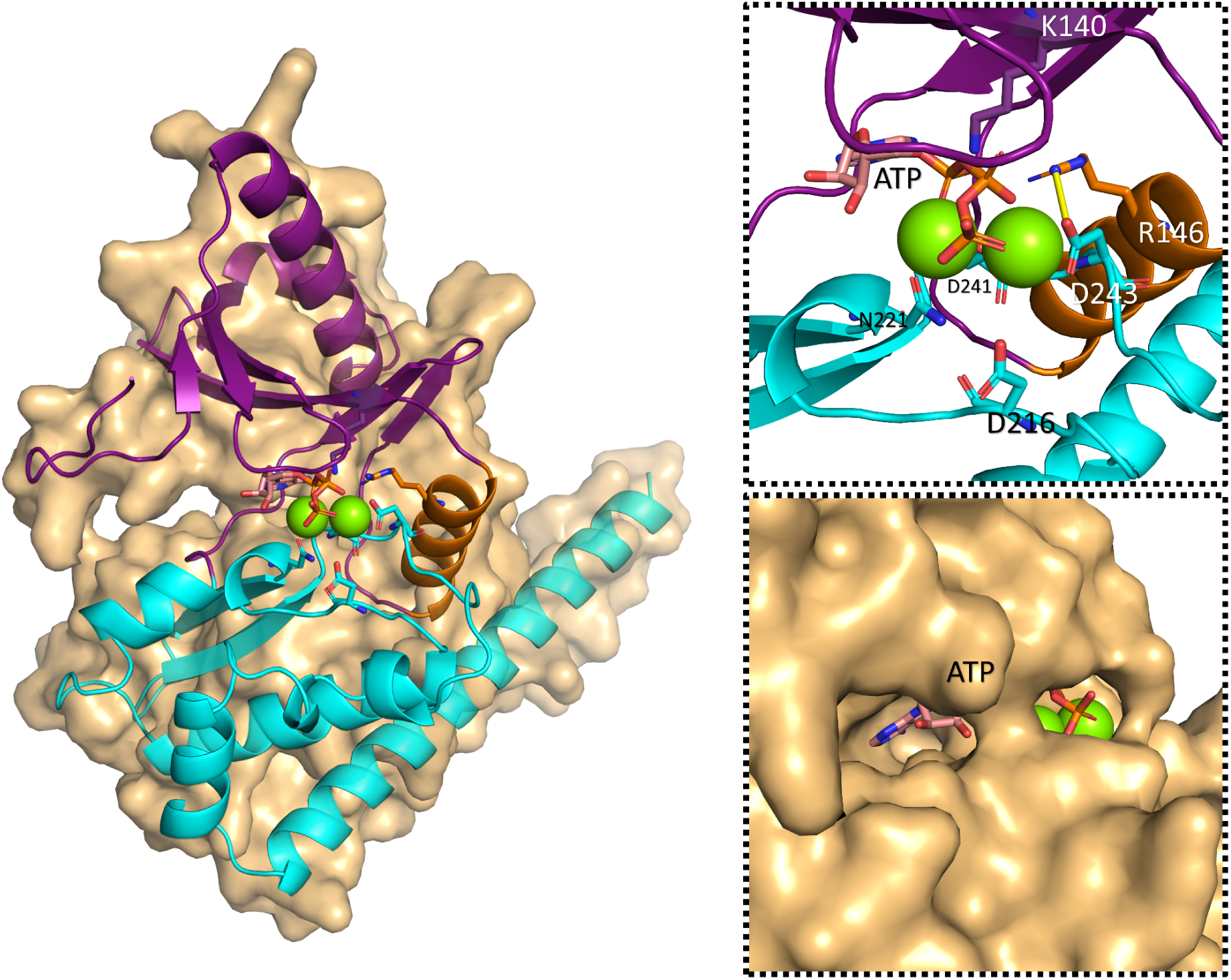
AlphaFold3 model of SEAE1. The overall structure is shown in a combined surface (light brown) and cartoon representation. The N-lobe is colored violet, the C-lobe cyan, and the αC-equivalent helix highlighted in orange. The binding pocket accommodates ATP (pink/orange sticks) and two divalent magnesium ions (green spheres) [27,35,36]. Zoomed-in view: the catalytic site showing key residues interacting with the ligands (K140, R146, D216, N221, D241, D243). The yellow line indicates a salt bridge (3.1 Å) between R146 and D243, this being structurally analogous to the conserved K–E ion pair which is typical of active protein kinases [27]. On the right is a surface representation showing the nucleotide-binding pocket occupied by ATP. The model confidence scores are pTM = 0.88 and ipTM = 0.96 (Suppl. Data 4A - SEAE1_models_AF3) [36].

**Figure 7.**
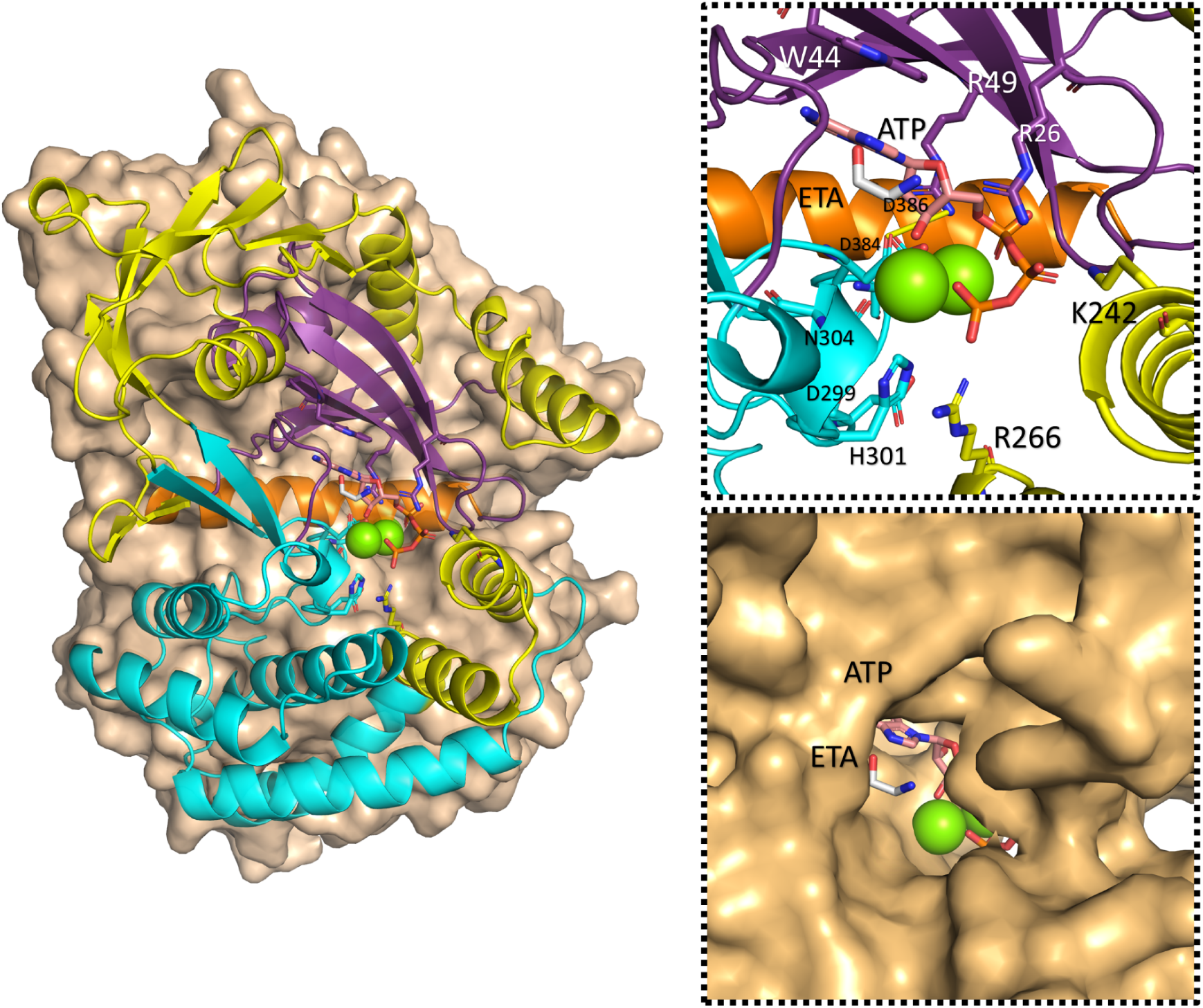
AlphaFold3 model of POGI1. The overall structure is shown in a combined surface (light brown) and cartoon representation. The N-lobe is colored violet, the C-lobe is cyan, the αC-equivalent helix is orange, and the inserts are yellow. The binding pocket accommodates adenosine triphosphate (ATP, pink/orange sticks), a putative substrate - ethanolamine particle (ETA, gray sticks), and two divalent magnesium ions (green spheres). Zoomed-in view of the catalytic site showing key amino acid residues interacting with the ligands (R26, W44, R49, K242, R266, D299, H301, N304, D384, D386). The yellow line denotes a salt bridge (4.5 Å) between R49 and D386. Bottom right: Surface representation highlighting the binding pocket occupied by the ATP and ETA. Model confidence scores: pTM = 0.95, ipTM = 0.96 (Suppl. Data 4B - POGI1_models_AF3) [36].

Analysis of the operon containing the kinase domain, along with effector prediction analysis (Suppl. Tables 5B and 3B), indicates that this protein is intracellular [42].

The operon organization is conserved in strains TDC60 and KCOM 3131. The operon (Supplementary Table 5B) includes XTP/dITP diphosphohydrolase, a small membrane protein, thymidine kinase (of the AAA/P-loop family), a putative PKL kinase, and 5′-methylthioadenosine/S-adenosylhomocysteine deaminase. This operon includes genes responsible for maintaining nucleotide pool quality (XTP/dITP hydrolase and thymidine kinase, which ensure the availability of normal DNA precursors) [62,63], and for the methionine cycle (MTA/SAH deaminase, which enables the recovery of methionine and adenine) [64,65]. The organization of these genes suggests that their functions are interdependent and may be co-activated in response to specific environmental conditions. *P. gingivalis* inhabits the periodontal environment, where it frequently encounters oxidative stress and nutrient deficiencies during host colonization [66,67].

*P. gingivalis* resides within a biofilm alongside other microorganisms, producing proteases and metabolites that alter the local microenvironment [68,69]. Efficient removal of S-adenosylhomocysteine (SAH) prevents this methylation inhibitor from accumulating, a factor which could affect the epigenetic control of virulence genes as well as the production of autoinducer-2 (AI-2) through the LuxS/SAM cycle. Although *P. gingivalis contain*s the luxS gene, AI-2 signaling appears to play atypical regulatory roles in this organism [64,70,71]. Additionally, efficient membrane metabolism via ethanolamine kinase may facilitate membrane remodeling, such as increased phosphatidylethanolamine levels or altered lipid content, which are associated with resistance to antimicrobial peptides and modulation of the host immune response in *P. gingivalis* [72–74].

The efficient functioning of these pathways likely enhances *P. gingivalis* ability to colonize the host and promote disease progression. Within the periodontal environment, the bacterium encounters heme limitation-heme is the primary iron–porphyrin source and is essential for P. gingivalis growth and virulence, along with oxidative stress and immune challenges. Survival is achieved through coordinated metabolic adaptations, including heme and iron scavenging, oxidative stress defenses, and outer membrane or lipid A remodeling. These adaptations support biofilm persistence, immune evasion through reduced TLR4 activation, and increased tolerance to cationic antimicrobials [73–79]. Membrane remodeling mediated by ethanolamine-modifying enzymes, such as increased phosphatidylethanolamine or altered lipid content, is associated with resistance to antimicrobial peptides and modulation of the host immune response in *P. gingivalis* [59,73,74,80–83].

The species-specific distribution and non-canonical active site geometry (R49–D386) distinguish POGI1 from human kinases. Bacterial MTA/SAH deaminase and XTP/dITPase in the surrounding operon likewise lack close human functional counterparts [84,85], making the operon an interesting candidate for future selectivity studies. We emphasize that this remains a bioinformatic observation: antimicrobial potential, target essentiality, and the proposed membrane-remodeling function would all require experimental validation before any therapeutic relevance can be assessed [86].

In summary, POGI1 is a *P. gingivalis*-specific kinase-like protein situated within a housekeeping operon linked to nucleotide quality control, methionine salvage and potentially membrane remodeling. The fact that it has a non-canonical active site structure and a limited phylogenetic distribution indicates that it has a specialized, pathogen-specific function. The catalytic activity of POGI1 and its physiological substrate remain undetermined. If POGI1 is confirmed as an ethanolamine or choline kinase, its activity could intersect with membrane lipid homeostasis and antimicrobial susceptibility, although this relationship is currently speculative. Definitive functional characterization will have to wait until biochemical tests have been carried out and genetic studies have directly linked POGI1 to membrane remodeling and virulence before its potential as a drug target can be assessed.

### Convergent evolution of the PKL scaffold in SrfA toxin

Analysis revealed that the SrfA protein contains a domain with remote structure similarity to PKL proteins (Suppl. Table 2). SrfA proteins of three organisms were examined: *Haemophilus parainfluenzae* (CBW15734.1, an upper respiratory tract commensal and opportunistic pathogen), *Xenorhabdus stockiae* HN_xs01 (QBB68691.1 (plasmid - XRZ01887.1)), and *Salmonella enterica* LT2 (NP_460552.1).

SrfA is a component of the three-part SrfABC toxin system, functioning as the effector protein. Along with SrfB and SrfC, it forms a functional toxic complex whose activity depends on the interaction of all three components [87,88].

In *S. enterica*, SrfABC was identified as part of the SsrB regulon. SsrB is a major response regulator that controls Salmonella Pathogenicity Island 2 (SPI-2), which is essential for the bacterium’s ability to survive inside host cells and for its virulence. Nevertheless, the specific biological function of SrfABC within this regulon has not been determined [89,90]. Experimental data regarding activity are available for *X. stockiae,* an entomopathogenic symbiont of *Steinernema nematodes*. In this organism, SrfABC functions as a tripartite toxin with injectable insecticidal activity against *Helicoverpa armigera* larvae and cytotoxicity toward insect midgut CF-203 cells [87,89]. The toxin was subsequently shown to induce apoptosis in mammalian cancer cell lines, with all three components capable of independent entry into cell but requiring the full complex for maximal activity [87].

Structural models generated by ColabFold (AlphaFold2) and AlphaFold3 indicate that SrfA serves as a platform to which SrfB binds first, followed by the association of SrfC to the preformed complex (Figure 8) [36,58].

**Figure 8.**
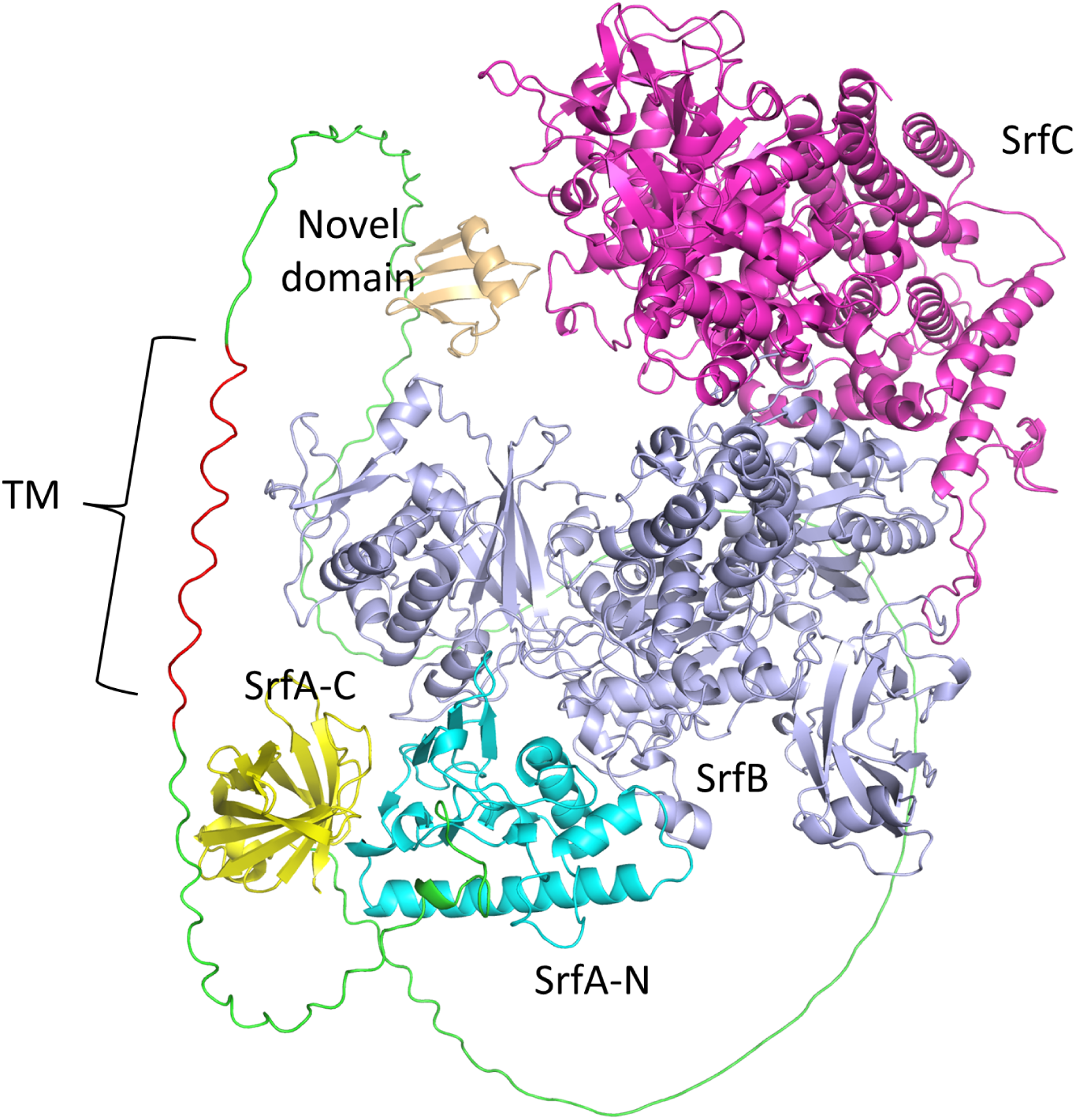
AlphaFold3 model of the Haemophilus parainfluenzae SrfABC toxin complex. The SrfA subunit is composed of multiple regions: the N-terminal domain (SrfA-N, cyan), a disordered region (green), a transmembrane helix (TM, red), a novel domain (ecru), and the C-terminal domain (SrfA-C, yellow). The interacting SrfB and SrfC proteins are depicted in marine blue and magenta, respectively. Model confidence scores for the full complex are pTM = 0.71 and ipTM = 0.66 (Suppl. Data 4C - SrfABC_models_AF3).

SrfA consists of three domains: an N-terminal kinase-like domain (SrfA-N), a transmembrane (TM) segment, and a C-terminal lipocalin-like domain (SrfA-C).

`BLAST analysis revealed that SrfA-N domains have homologs predominantly` in Gram-negative bacteria (E-value = 1e−4). Many of these organisms are human pathogens, including *Pasteurella multocida, Yersinia pestis, Serratia marcescens, and Pseudomonas aeruginosa. H. parainfluenzae* is a relatively commensal gram-negative microorganism that inhabits the upper respiratory tract, but can also act as an opportunistic pathogen [91]. The SrfA-N domain (respectively 1-165, 1-143 (1-132), 1-131) lacks all key catalytic motifs characteristic of protein kinases (“VAIK,” “HRD,” “DFG”) and does not bind ATP according to AlphaFold 3 (Suppl. S2A) [27]. The sequence logo (Suppl. Data 2) shows that the conserved residues in this domain are entirely different from those conserved in canonical kinases. Such a pattern indicates that, although the domain adopts a PKL fold, it is not part of the kinase lineage. On the contrary, it is a member of a kinase-like family which very likely evolved through convergent evolution, involving the independent adoption of a PKL structural scaffold. At the same time, the sequence constraints and functional determinants underlying it have diverged entirely from those found in true kinases.

The subsequent transmembrane helices (positions 208-227, 220-239 (209-230), and 232-251, respectively) likely anchor the TM protein within the bacterial inner membrane bilayer. Notably, AF2/3 fails to model this element accurately. DeepTMHMM identifies a transmembrane region in all SrfA proteins (Suppl. Table S5C-E, Suppl. Table S6A) [92–94]. Furthermore, CCTOP server (Consensus Constrained TOPology prediction) was employed to validate the more accurate predictions in SrfA from *X. stockiae, H. parainfluenzae*, and *S. enterica* (Suppl. Table S6B) [95].

The third domain at the C-terminus (406-525, 344-464 (333-453), 313-439) adopts a typical lipocalin β-barrel structure, resembling an avidin-like domain [96]. The sequence logo of the “SrfA-C” family (Suppl. Data 4C - SrfA-C) demonstrates the great majority of sequences retain the characteristic motif of the lipocalin subfamily, FLNGxW. This motif represents a modified version of the initial lipocalin motif GxW, followed by partially conserved block TGKP and, in the second half of the domain, a CxDGS motif containing the first of four conserved cysteines [96,97]. Approximately 89.5% (1595/1782) of the analyzed C-terminal domain sequences contain four cysteine residues positioned to enable the formation of two disulfide bridges. As disulfide bridges predominantly form in the oxidizing environment of the periplasm and extracellular spaces, where disulfide bond (Dsb) systems function, this cysteine arrangement supports the periplasmic localization of the domain [97–99]. Lipocalins are involved in OMV transport [100].

Genomic analysis indicates that in both *H. parainfluenzae* and *X. stockiae*, the *srfABC* operon is flanked by a conserved gene cassette encoding a Lol-like ATP-binding cassette (ABC) transporter module (comprising ATPase and permease), von Willebrand factor A (vWA) domain-containing proteins, and a SUMF1/EgtB/PvdO family nonheme iron enzyme. This enzyme, a formylglycine-generating enzyme, is potentially involved in periplasmic post-translational modification of SrfA-C [87–89,101–105]. HHpred analysis (Suppl. Table S5C-E) of the permease components (CBW15740.1 in *H. parainfluenzae*, XRZ02085.1 in *X. stockiae*) demonstrates dual similarity to both LolC and LolE subunits, which is characteristic of LolF-type permeases that homodimerize to perform combined chaperone and substrate recognition functions. Each permease associates with an ABC ATPase (CBW15739.1/XRZ02097.1), forming a simplified two-gene Lol system that is functionally analogous to the canonical three-gene LolCDE complex [29,106,107]. Both organisms also possess canonical chromosomal LolCDE operons which are involved in the transport of housekeeping lipoproteins (CBW15551.1, CBW15552.1, CBW15553.1 and XRZ04735.1, XRZ04734.1, XRZ04733.1), indicating that the SrfABC-associated modules are specialised for toxin processing rather than serving as a replacement for the essential machinery.

In contrast, *S. enterica* lacks this specialized module entirely, even though it does encode a homologous SrfA.The fact that it lacks this module indicates the possibility of other export mechanisms or a lower demand for secretion, potentially reflecting distinct pathogenic lifestyles. The *S. enterica* SrfABC locus is instead flanked by genes typical of horizontally acquired pathogenicity-associated regions, including a GhoT/OrtT family type V toxin–antitoxin module [108,109]. This arrangement, which is specific to the species, lends support to the idea that Lol modules play a functionally important role in optimising the export of SrfABC, rather than simply showing a coincidental co-localisation [29,110].

The absence of Sec/Tat signal peptides (Suppl. Table S6C [44,100,111–114]) and the presence of a stable transmembrane helix exclude transport by T1-6SS, which require cytoplasmic, soluble substrates that are not membrane-anchored. The “secreted effector” annotation in CDD (NF040486) originates from the Salmonella SPI-2 context [115,116], rather than from protein localization predictions. Notably, in Salmonella, the SrfABC operon is not co-expressed with SPI-2 genes, its products are not translocated via T3SS [89], and SrfA maintains identical bitopic topology. These observations collectively undermine the validity of the “secreted effector” designation.

Outer membrane vesicles (OMVs) are known to act as carriers for the export of toxins, virulence factors, and membrane-associated proteins in Gram-negative bacteria [117,118]. Beyond classical OMVs, outer–inner membrane vesicles (O-IMVs) can transport proteins embedded within their native membrane environment, including bitopic membrane proteins [100]. The production of OMVs is common among various Gram-negative bacteria, including those in the *Pasteurellaceae* family. In *H. parainfluenzae*, outer membrane-derived vesicles (40–70 nm) are formed by budding from the cell envelope, and vesicle-associated DNA-binding activity has been connected with genetic transformation mechanisms [119–121]. The delivery of membrane-anchored proteins is made possible by vesicle-mediated export, since the periplasmic folding conditions, such as the formation of disulfide bonds, which are necessary for protein stability and activity, are maintained [98].

The results point to OMV-mediated secretion in the export of SrfABC in *H. parainfluenzae and X. stockiae*, and show differences between species in the structure of their export systems in *S. enterica*. The comparative study of these pathogens, which occupy different ecological niches, provides a basis for clarifying uncertain toxin annotations and underscores the functional diversity of OMV-dependent virulence mechanisms. The fact that SrfA-N has been assigned to the PKL superfamily is based on structural similarity of the fold, not on any demonstrated catalytic activity or substrate binding. Since the catalytic motifs have diverged, the protein is probably not functioning as a kinase, and if it has any function left beyond the scaffolding role suggested here, that function may involve substrates and a type of chemistry other than phosphoryl transfer. It cannot be ruled out on the basis of structural evidence alone that convergence on a common fold has occurred from unrelated ancestral origins. Resolving this point will require experimental investigation, and the family therefore presents an attractive target for such studies.

## Conclusions

Through bioinformatic analysis of oral microbiome bacteria, we identified 20 novel PKL families. We projected the entire superfamily onto a comprehensive map of PKL structure and sequence families to validate their relationships. By integrating AI-based structure prediction with systematic taxonomic and genomic analyses, we identified PKL-containing proteins exhibiting distinct evolutionary distributions and functional adaptations. Three of these proteins are described in detail in Results.

The SEAE1 protein kinase family is found in a wide variety of bacterial phyla and has been identified in 7,867 homologs. It is predicted to have kinase activity as part of a PAP2-PIK metabolic module. The fact that it is widely distributed taxonomically and that its operonic structure is conserved suggests that it plays important fundamental roles in lipid signaling pathways.

POGI1 exhibits high phylogenetic specialization, being almost exclusively restricted to *Porphyromonas gingivalis* strains. Structural modeling supports its role as bacterial PKL-fold ethanolamine kinase. The POGI1 gene is located within an operon involved in coordinating nucleotide quality control, methionine salvage, and membrane remodeling, all of which are important adaptations for survival in the periodontal environment.

SrfA-N is an example of convergent evolution of the PKL scaffold for non-catalytic functions and is widely found amongst Gram-negative pathogens as a toxin platform. Analysis based on systematic exclusion has shown that export via outer membrane vesicles is the primary secretion mechanism, correcting earlier annotations regarding the T3SS and revealing sophisticated vesicle-dependent virulence mechanisms.

Together, these three families show that PKL domains have evolved along several distinct paths: enzymatic specialization (SEAE1), adaptation to specific pathogens (POGI1), and structural co-option (SrfA-N). The range of species-specific distribution patterns, from widely conserved to highly restricted, illustrates the role of protein-fold versatility in enabling bacteria to adapt to diverse environmental and host-related lifestyles.

A primary limitation of this study is its reliance on computational predictions without experimental validation. Nevertheless, the distinct structural features, taxonomic distributions, and genomic surroundings provide strong support for the suggested functional assignments. These proteins do constitute promising targets for use as antimicrobials, especially POGI1, since it is distributed specifically among pathogens, and SEAE1 because it has an essential metabolic function.

Our systematic approach, which combines sequence and structure prediction with mechanistic analysis, establishes a framework for discovering functionally significant protein families within complex microbial communities. Further progress in research on the oral microbiome will likely lead to the discovery of similar PKL-mediated adaptations in other bacterial lineages, showing that continuous evolutionary innovation occurs within existing protein folds.

Novel PKL families, including sequences, alignments, HMM models, and 3D PKL domains, are available on (zenodo link). In the future, these resources will supplement our database (http://bioinfo.sggw.edu.pl/kintaro/).

## Materials and methods

### Search strategy

Protein sequences from HOMD v9.15a (5,1mln) were clustered at 90%, 60%, and 30% sequence identity using CD-HIT (PSI-CD-HIT (default) at 30% sequence identity) [2,122–124]. Sequences were divided into 300-amino-acid fragments with 50-amino-acid overlap, and fragments shorter than 150 aa were discarded [125]. As a pre-filtering step prior to structural modeling, fragments matching known protein domains were removed using RPS-BLAST against the NCBI Conserved Domain Database (CDD 3.19, E-value = 1e-3) [116] and HMMER [126] against Pfam 35 [127] and KINtaro [8] ECOD 282 databases (E-value = 1e-4). The remaining sequences had structures modeled using ESMFold (64544) [128]. As a structural pre-filter, predicted structures were compared against a curated set of known PKL structures from the ECOD40 282 (only ECOD40 entries with a single, contiguous polypeptide chain were used) [126] and KINtaro [8] databases using TM-align version 20220227 [28] (significance threshold: TM-score = 0.4). Candidate PKL families were further validated using LogoSS, HHpred (PDB70_mmCIF70 [129], ECOD_F70 [126], Pfam-A, COG_KOG, NCBI_Conserved_Domains(CD)) [29], FATCAT (PDB90) [31], and DALI (PDB90) [30].

### Sequence data

Protein sequence data for novel kinase-like families were provided from HOMD Genomic Reference Sequences version 9.15a [122].

### Clustering protein sequences

Due to the large size of the sequence data, the sequences were clustered by sequence identity using the CD-HIT suite. Three clustering thresholds were used: 90%, 60% and 30% sequence identity. This reduces the load on the processor and graphics card [123,124].

### Splitting sequences

Sequences were split into 300 aa long segments with an overlap of 50 aa [125].

### Discarding known domains

The sequences were discarded using RPS-BLAST (CDD, E-value = 1e−3) [32,116] and HMMER (Pfam and KINtaro HMM, E-value = 1e−4 [8,127,130]).

### Rejecting short sequences

We used in-house scripts to reject short sequences under 150 aa [125].

### Genomes/Assemblies

Genomes/Assemblies were obtained from NCBI (Suppl. Data 4 - Genomes) [131].

### Remote homology detection and structure modeling

For distant similarity detection to PKL families, we used the TM-align algorithm [28], our ESMfold models [128] vs PKL proteins from KINtaro [8] and ECOD40 structure set [126]. To validate our hypotheses, we used FATCAT (PDB90) [31,129], Dali (PDB90) [30,129] and HHpred [67] (PDB70 [129], ECOD F70 [126], SCOPe70 [132], Pfam [127]). All with standard parameters. Novel PKL-like structures (and known in Suppl. Table S4) were modeled with the use of ColabFold (AlphaFold2) (Suppl. Table S2: 24 cycles, Suppl. Table S4: default) [58], AlphaFold3 server (Suppl. Table S2 and 4: default) [36] and ESMFold (Suppl. Table S2 (additional analysis and S4: 24 cycles) [128]. The best models have been selected. Structural comparisons were performed using the FATCAT and Dali servers (Suppl. Table S2) [30,31].

### AlphaFold3 docking

All docking runs used 20 random seeds per condition, and models were ranked by ipTM score, the top-ranked model per condition was retained for structural analysis [36].

SEAE1 was modeled with ATP (ChEBI:30616). Two magnesium ions (ChEBI:18420) were included [61].

For POGI1, three ligand conditions were evaluated: 1. ATP alone, 2. ATP with the ethanolamine in the neutral form (NCCO, ChEBI:16000), 3. ATP with the ethanolamine in the protonated form ([NH3+]CCO, ChEBI:58243). Two magnesium ions (ChEBI:18420) were included in all conditions [36,61].

### Previously characterized PKL families histogram chart

The HOMD v9.15a proteins were searched using hmmsearch (E = 1e-4) against the 71 KINtaro family profiles. Each protein was assigned to the family associated with the profile that gave the highest full-sequence bitscore. The first analysis revealed 36 families; of these, nine families with best-hit bitscores below 50 were further assessed based on alignment and catalytic-motif content rather than discarded solely because of the bitscore threshold. Six families (Pan3_PK, Kinase-like, Pkinase_fungal, UL97, TCAD9, Lqui_0983) were excluded, leaving 30 reported families and 10,615 proteins. Three families with weak best hits (Lspi_2187, HopBF1, Act-Frag_cataly) are listed as provisional in Suppl. Data Table 1. The number of proteins may be affected by how strains are sampled in HOMD; hence, the HMT taxon counts provide a more conservative estimate of the distribution.

### Comparison of PKL family databases

Overlap between the 20 HOMD PKL families identified here, the KINtaro database (n=71) [8], and Pfam protein kinase clan CL0016 (n=48), InterPro, accessed 26.08.05) was assessed by matching KINtaro entries to Pfam accessions [127,133]. Three HOMD families previously deposited in KINtaro as preliminary results were assigned to the KINtaro ∩ HOMD overlap. Results were visualized as a Venn diagram (Figure 2) [8] Pfam clan CL0016 contains 48 accessions, which are consolidated into 46 families according to the family definitions provided by the KINtaro database (PF00069/PF07714 = PKLF000033; PF07804/PF20613 = PKLF000019).

### Taxonomic distribution analysis of PKL homologs

The phylogenetic breadth of each PKL family was assessed using BLASTp searches against the nr database (E-value = 1e−4, NR 2025), excluding sequences shorter than 100 amino acids, for all 20 representative sequences [32,125]. Taxonomic phylum assignments were resolved using the NCBI Taxonomy database (ete3 v3.1.3) [125,134,135], archaeal hits were excluded. The number of unique organisms per family per phylum was visualized as a log-scale dot plot (Figure 4).

### Generating a sequence logo with a secondary structure annotation - LogoSS (prototype)

Sequence conservation was visualized as an information logo with a secondary structure overlay using a custom Python pipeline (LogoSS, Suppl. Data 2). This pipeline integrates structure prediction [58], multiple sequence alignment (MSA) analysis [136], and the Logomaker library [137].

Protein structure prediction was performed using ColabFold with 24 recycles, resulting in both a PDB structural model and a multiple sequence alignment (MSA) generated by MMseqs2 [32,136]. Homologs with coverage below a specified threshold (default 0%) were optionally excluded. The MSA was then simplified by removing insertions and query-gap positions, producing sequences of the same length. Secondary structure was determined from the model using DSSP [138].

The sequence logo was generated from information content (bits) calculated from amino acid frequencies at each alignment position, excluding gaps. Amino acids were colored according to a functional scheme: green for polar (G, S, T, Y, C, Q, N), blue for basic (K, R, H), red for acidic (D, E), and black for hydrophobic and other amino acids. Information values below the noise threshold (default 0.05 bits) were set to zero. A secondary structure bar is displayed below each segment of the logo (default 60 residues): orange for helices, blue for β-sheets, and grey for loops or coils.

For pseudokinases, the characteristically diffuse sequence logos are consistent with the expected relaxation of catalytic constraints [10]. Since pseudokinases retain the PKL fold for regulatory or scaffolding roles, there is no need for strict conservation of the active-site residues, and the observed sequence variability is therefore probably due to an underlying biological signal rather than merely to methodological limitations.

The current implementation generates structures and MSAs internally via ColabFold; future versions of LogoSS will support user-supplied PDB structures/models, secondary-structure annotations, and sequence alignments.

### Protein Structure Visualization

Protein models were visualized using PyMOL [35].

### Visual clustering of families (analysis of sequence similarity relations between families)

To visualize clusters of protein kinase families, the sequence clustering we used the CLANS [33] algorithm with the BLOSUM62 scoring matrix [139] and extraction of BLAST [32] hits up to an E-value of 1. The set of sequences was adopted from our previous work [9]. Novel families have been added. All sequences have been modeled (Suppl. Table 4) and experimental structures have been added. Those sequences/structures that deviate significantly from the groups in the graph have been removed.

### Visual clustering of families (analysis of structural similarity relations between families)

To visualize the structure/model similarity network of PKL families we used the NISARA script from our previous publication [34]. To visualize similarities between PKL protein structures, we used a TM-score cutoff threshold of 0.4 [28,33].

### Novel PKL families

`Homologs were collected using BLAST (E-value = 1e−4, NR 2025)` [32,131], sequences shorter than 100aa were rejected [125], rest were clustered by CD-HIT (50% id, 100% if family sequences < 10) [123,124], sequences were aligned with ClustalO [140]. Full-length sequences were retrieved by ID from domains [125,131].

Newly identified PKL families were designated using a five-character code comprising the first two letters of the genus and the first two letters of the species of the NCBI representative organism, followed by a numeric discovery-order suffix (e.g., SEAE1 - first novel family identified in *Segetibacter aerophilus*). Where species-level assignment was unavailable, the HOMD oral taxon number replaced the species component (e.g., AT416 for *Atopobium sp.* oral taxon 416). Family designations reflect the NCBI representative organism used for domain annotation and homology searches; original HOMD source genomes are documented in Suppl. Table S1.

Two exceptions to this convention apply. The designation SrfA-N is maintained in accordance with established literature, as it specifically refers to the N-terminal domain of the SrfA protein and avoids confusion with previous studies. LACHF and LACHP were named prior to adoption of the current convention, using a four-letter genus abbreviation (LACH - Lachnoanaerobaculum) followed by a single functional descriptor: F denoting the presence of an additional phosphatase domain, and P denoting pseudokinase character. These designations are retained for consistency with the KINtaro database [8] and our earlier study [26].

Underscores in family names were replaced with hyphens during computational processing (e.g., Choline_kinase → Choline-kinase) to avoid conflicts with the underscore field separator used in sequence identifiers (FAMILY_accession_coordinates). Both forms denote the same family.

### Substrates of secretion systems

Substrates of secretion systems were predicted with use of SignalP-6.0 - fast mode (2021) [44], BastionX - fast mode (2021) [141], T3SEpp (2020) [142], T4SEpp (2024) [143], T6CNN (2025) [144]. Homologs were identified by BLASTp (E-value = 1e−4, NR 2023) [32,131]. The retrieved domain sequences were clustered using CD-HIT at 50% sequence identity to reduce redundancy [123,124]. Full-length protein sequences corresponding to cluster representatives were then retrieved by identifier and subjected to effector function annotation (Suppl. Data 4 - Homologs).

In the BastionX dataset, the letter “X” was removed from the sequences. he BastionX threshold was set to 0.8.

Peptide signal of proteins in operons (Suppl. Table 5) were predicted with SignalP-6.0 - fast mode (2021) [44].

### Transmembrane helices

Transmembrane topology of novel kinase families (Suppl. Table 1E) was predicted with DeepTMHMM [93].

Transmembrane topology of operons (Suppl. Table 5) was predicted with DeepTMHMM [93].

Transmembrane topology of all SrfA homologs (Suppl. Data 4C Table 1) was inferred using three independent predictors: Phobius [94], TMHMM [92], DeepTMHMM [93]. In addition to the four homologs discussed here (CBW15734.1, QBB68691.1, XRZ01887.1, NP_460552.1), transmembrane topology was independently verified using CCTOP [95], confirming a membrane anchor in all four proteins. homologs were identified by BLASTp (E-value = 1e−4, NR 2023 [32,131]). The retrieved domain sequences were clustered using CD-HIT [123,124] at 50% sequence identity to reduce redundancy. Full-length protein sequences corresponding to cluster representatives were then retrieved by identifier and subjected to transmembrane topology prediction (Suppl. Data 4 - Homologs).

### Operon prediction

For operon predictions we used Operon-mapper web service [42].

## Supporting information

Suppl. Logo

## Acknowledgements

The authors thank Dr Krzysztof Pawłowski for consultations and critical reading of the manuscript. The research and data collection were supported by the Polish National Science Centre under grant no. 2019/35/N/NZ2/02844 awarded to M.G.

## Use of AI declaration

A large language model (Claude - Fable 5, Anthropic) was used to draft analysis scripts (Figure 1, 3, 4); all code was verified against the primary search output and all interpretations are the authors. The authors take full responsibility for the content of this manuscript.

## Author Contributions

Performed the experiments and analyzed the data: M.G, M.K, W.Ś, Z.J.. Prepared figures and/or tables: M.G, M.K. Wrote the original draft: M.G. Authored and reviewed drafts of the paper: M.G., M.K. Conceived and designed the experiments: M.G. All the authors approved the final draft.

## Data Availability

Sets of sequences, PDB of HOMD PKL families, scripts are available from zenodo: 10.5281/zenodo.2283456510.5281/zenodo.22834565

LogoSS: 10.5281/zenodo.22832949

And on an online database at http://bioinfo.sggw.edu.pl/kintaro/.

