## Supplementary material for "Systematic discovery of protein kinase-like domains reveals diverse evolutionary strategies in the human oral microbiome": Suppl. Logo

### Logo with secondary structure

**Legend:**

**K72**

**E91**

**D166**

**D184**

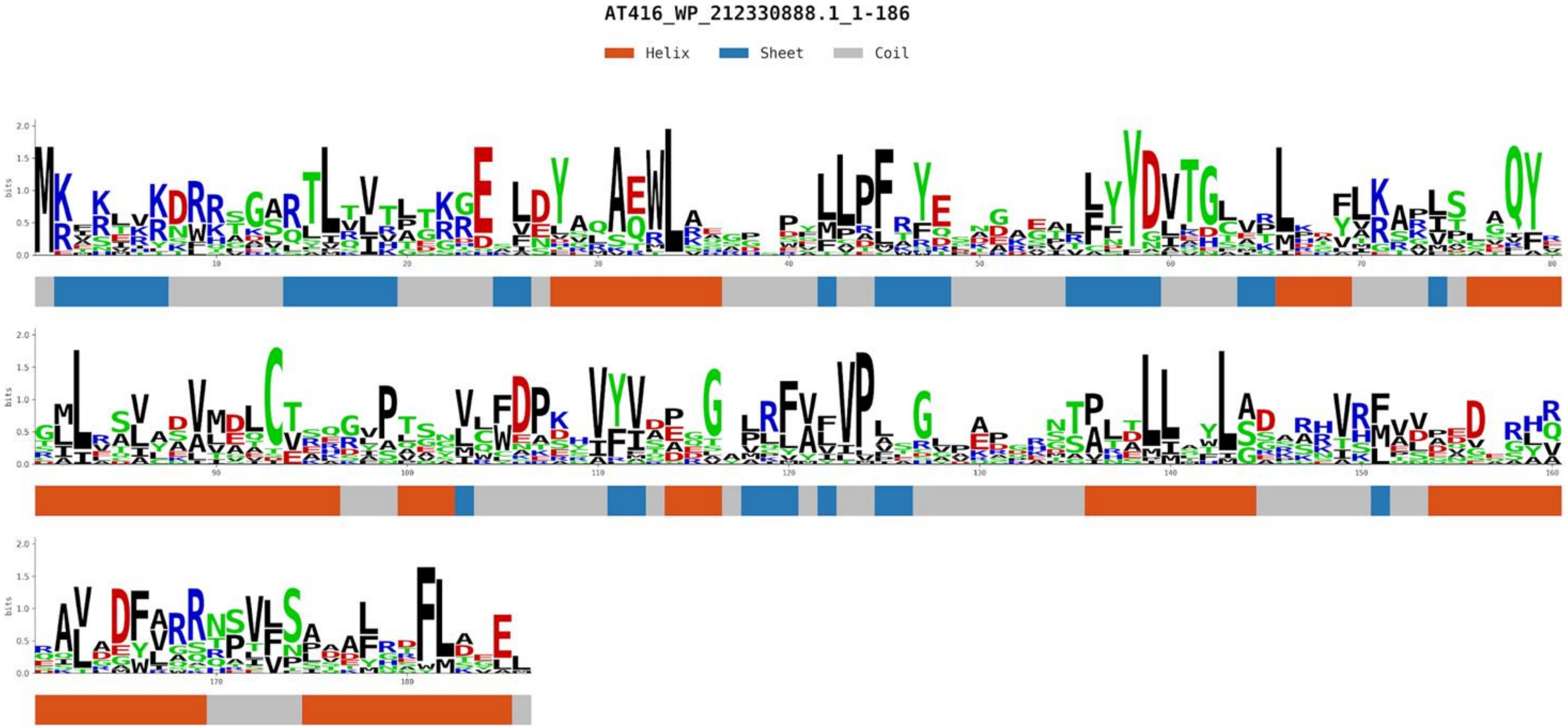


Only PKL fold recognized

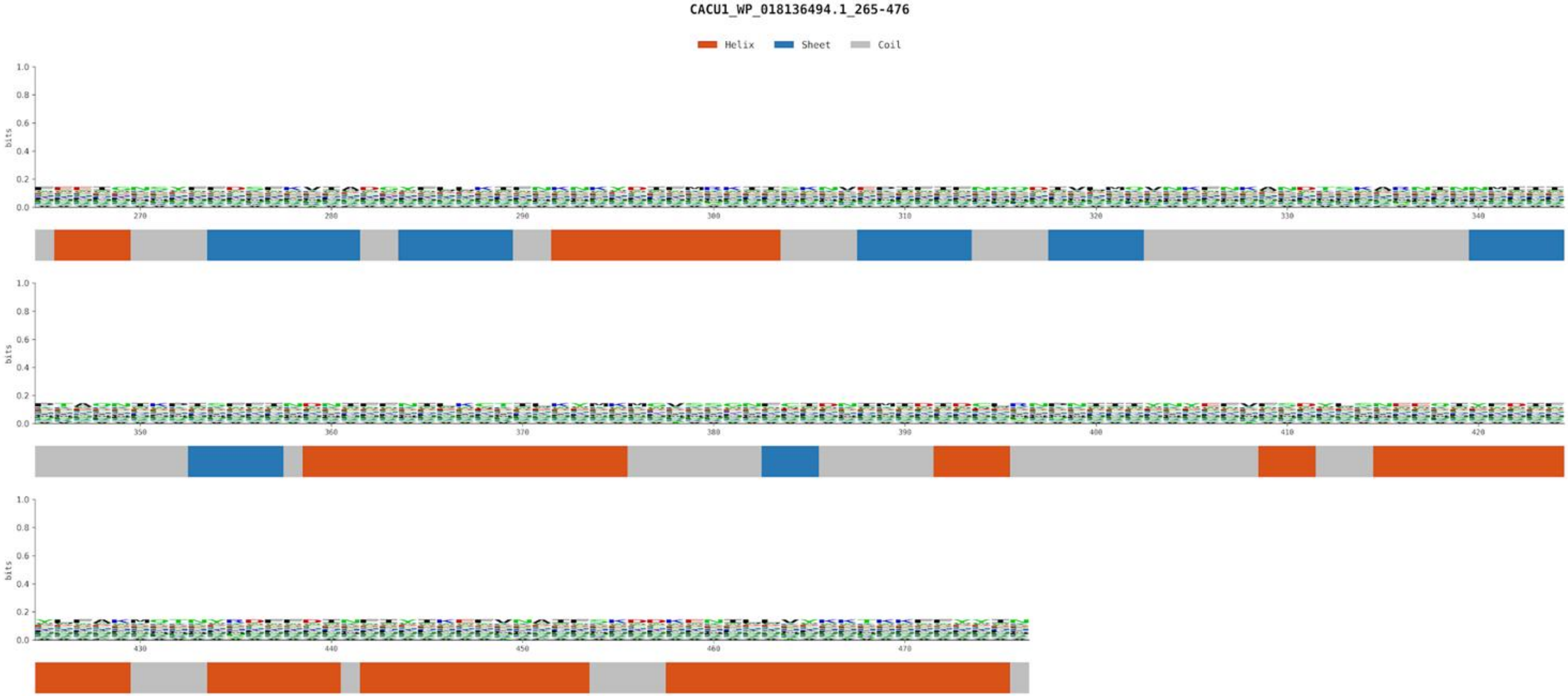


Only PKL fold recognized

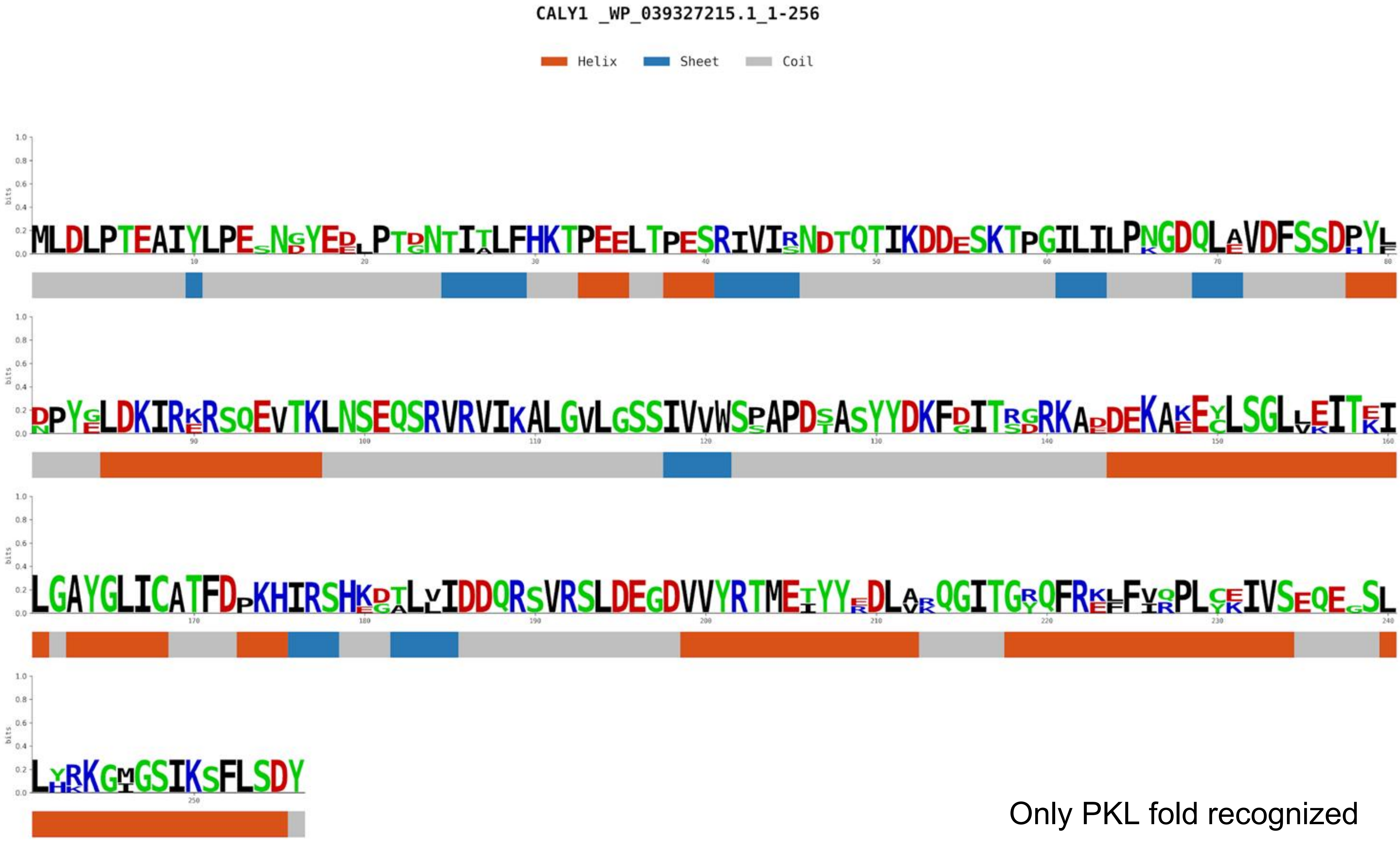


Only PKL fold recognized

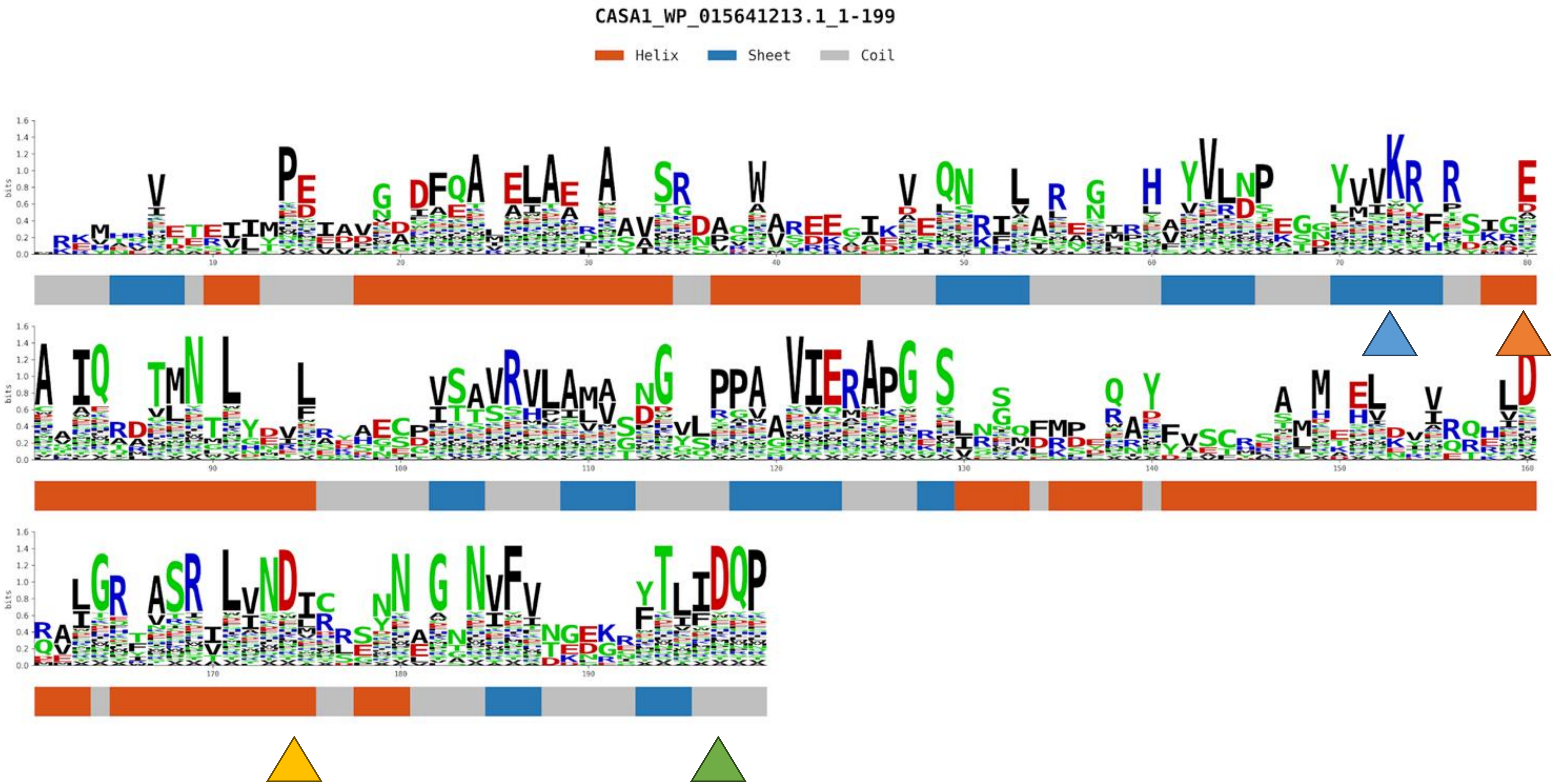
CASA1_WP_015641213.1_1-199
Helix
Sheet
Coil
bits

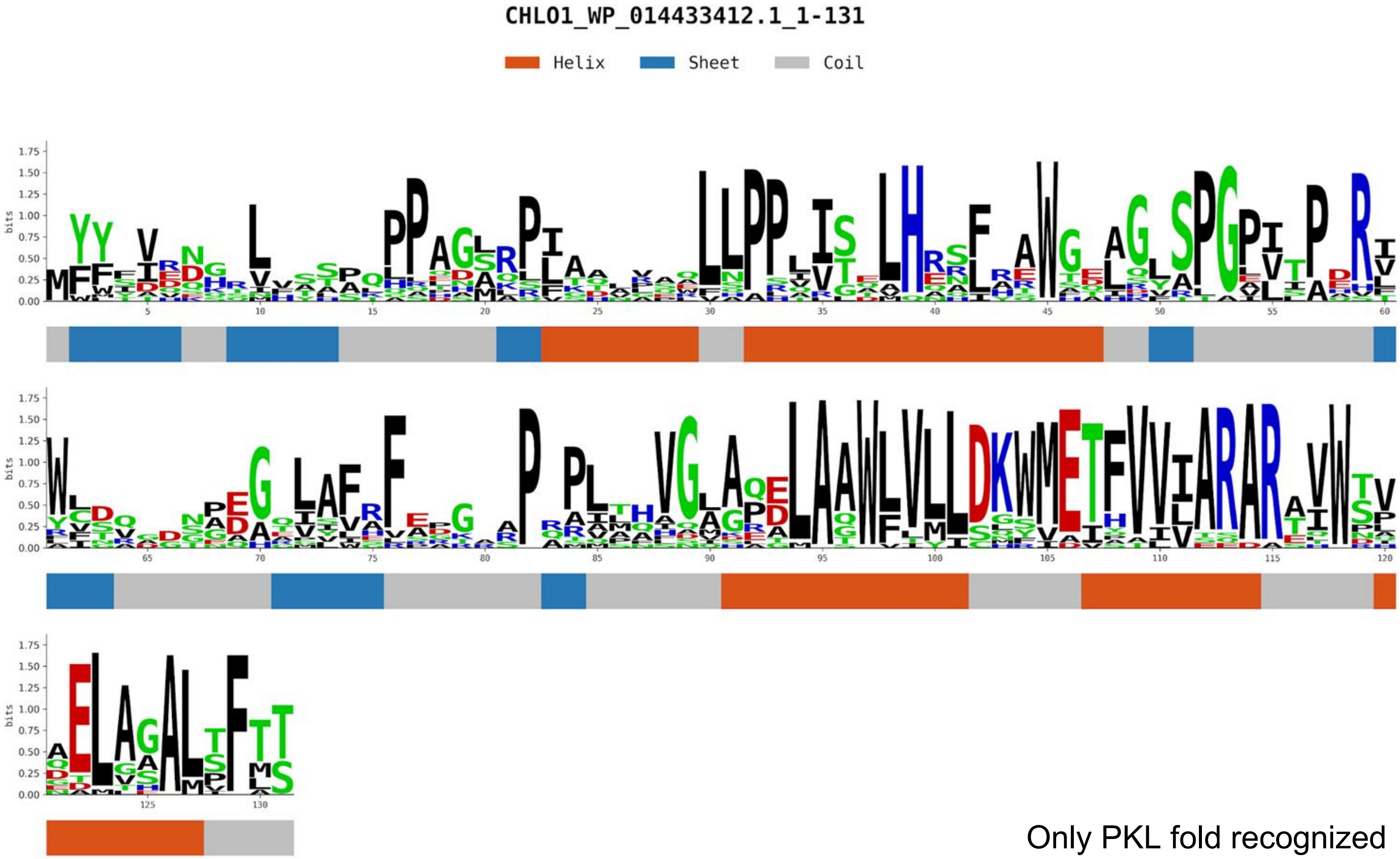


Only PKL fold recognized

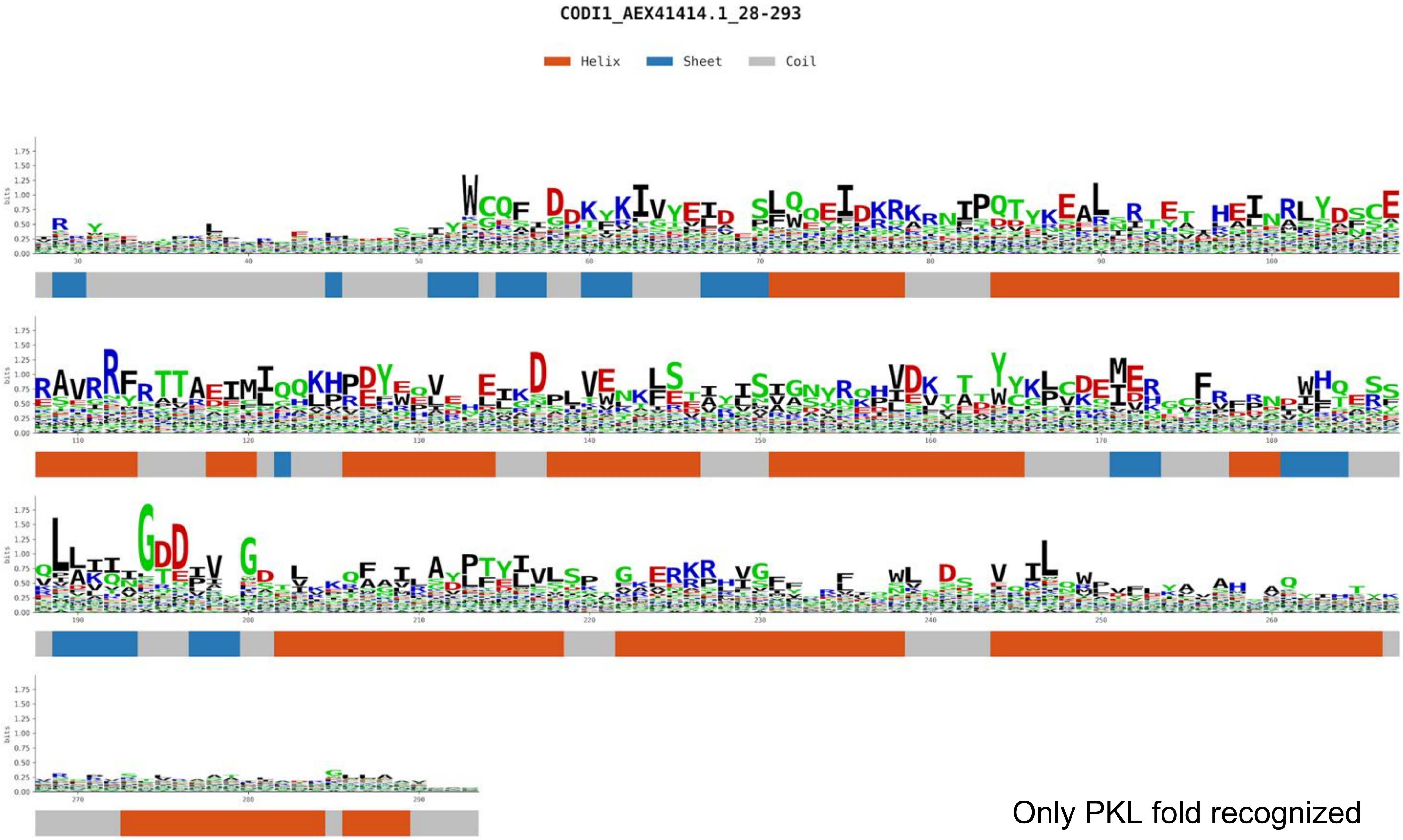


Only PKL fold recognized

CUGI1_WP_261667374.1_997-1180

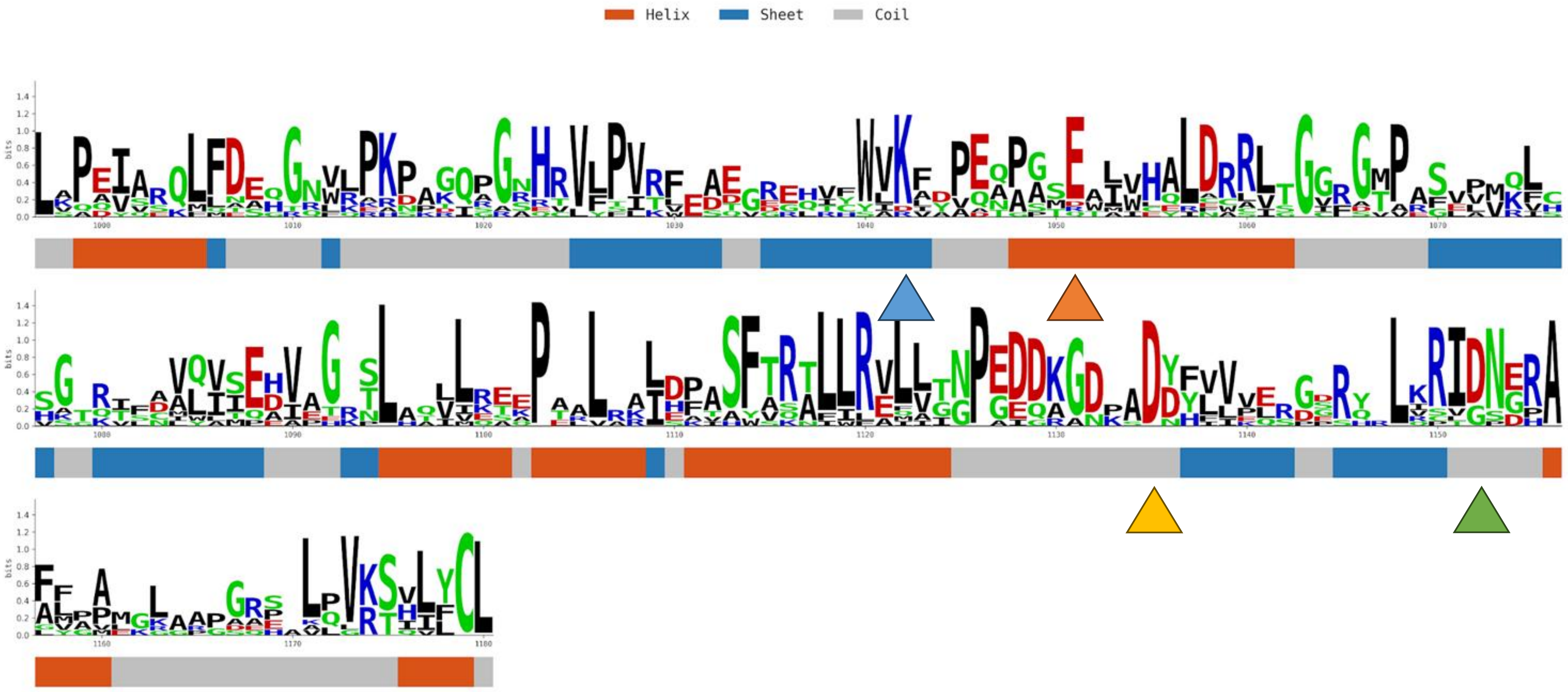

IGAL1_WP_014561518.1_1078-1263

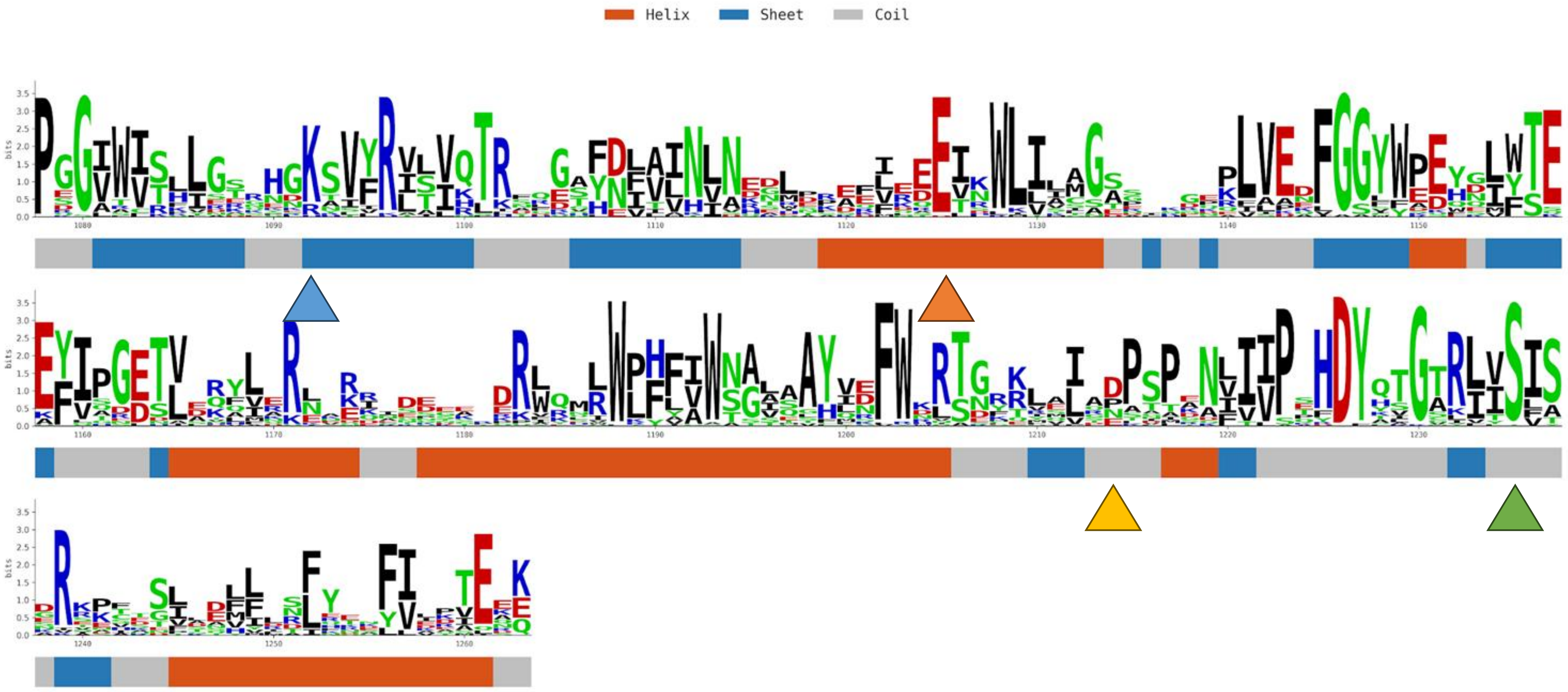

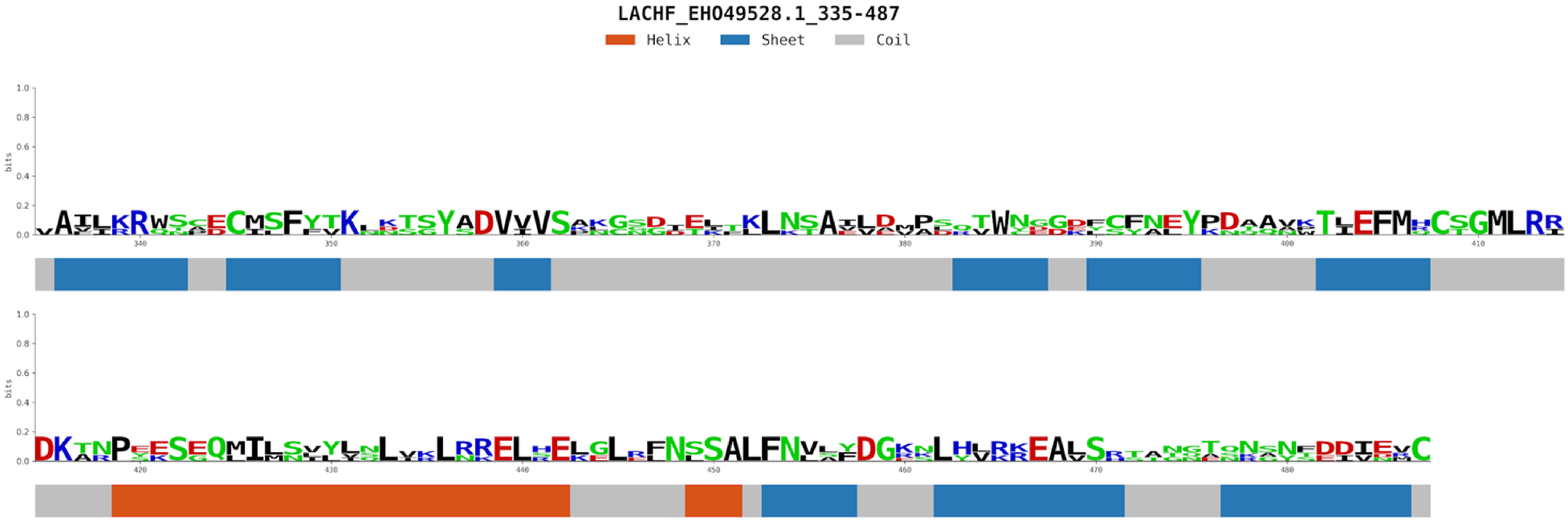


Only PKL fold recognized

LACHP_EHO49530.1_34-305

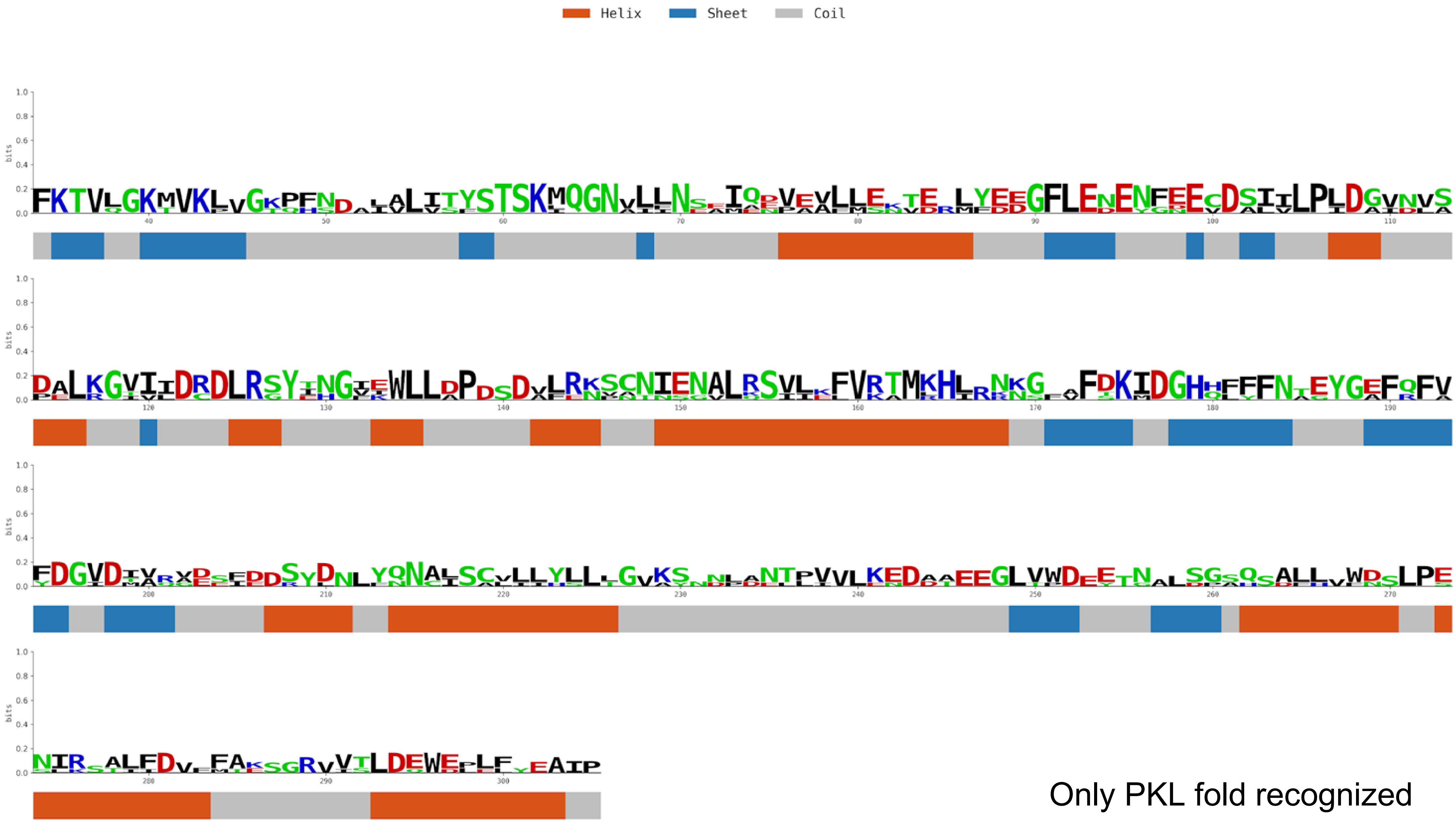


Only PKL fold recognized

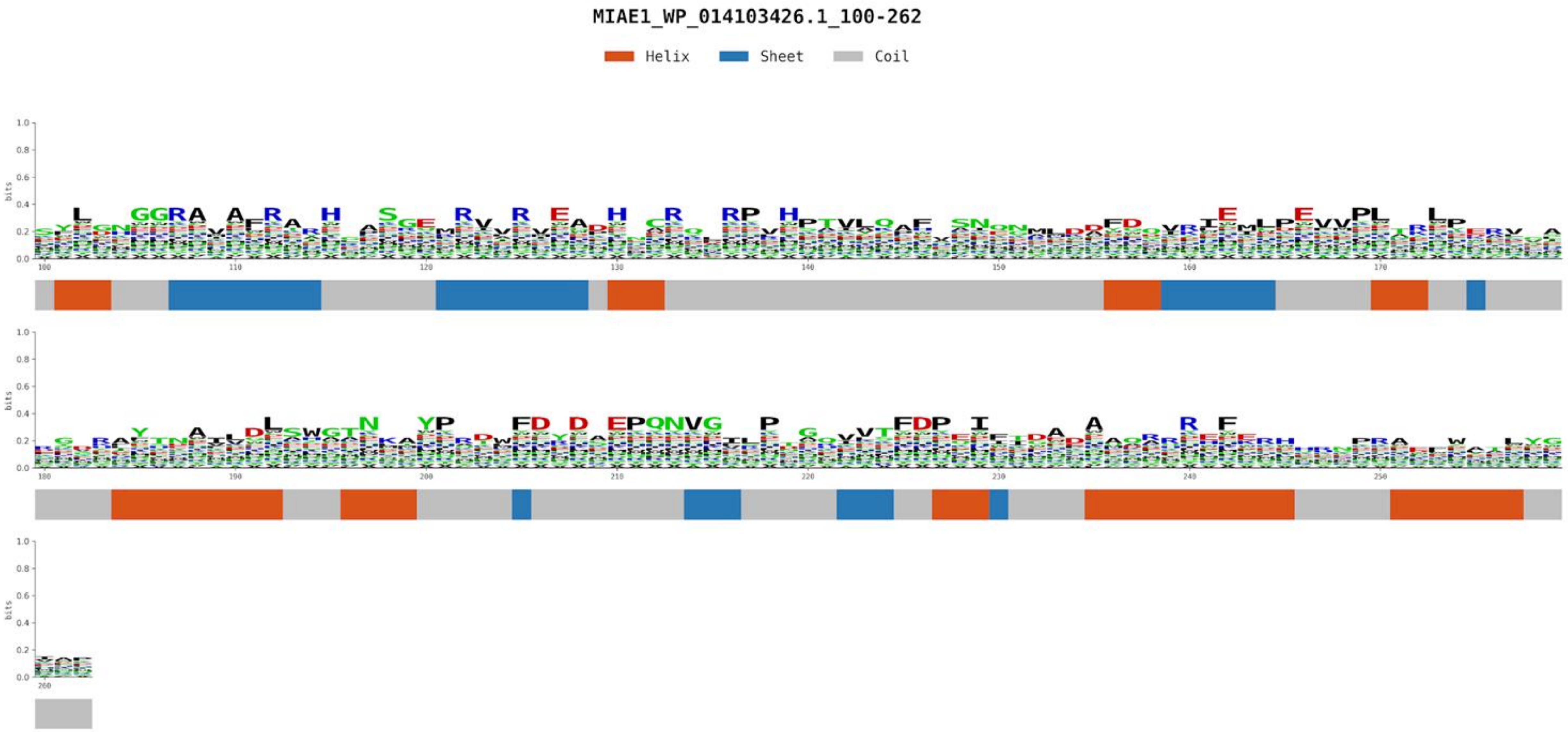


Only PKL fold recognized

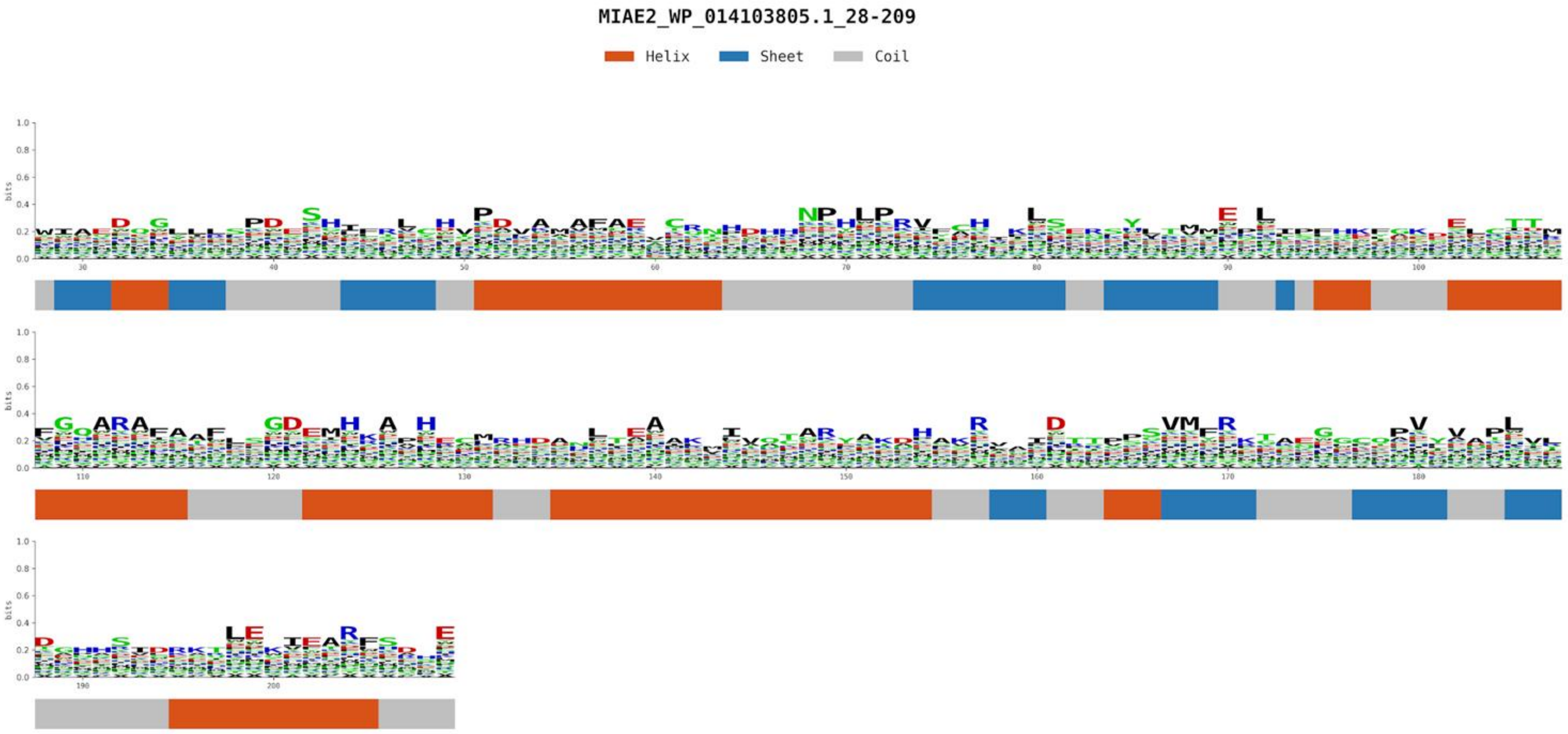


Only PKL fold recognized

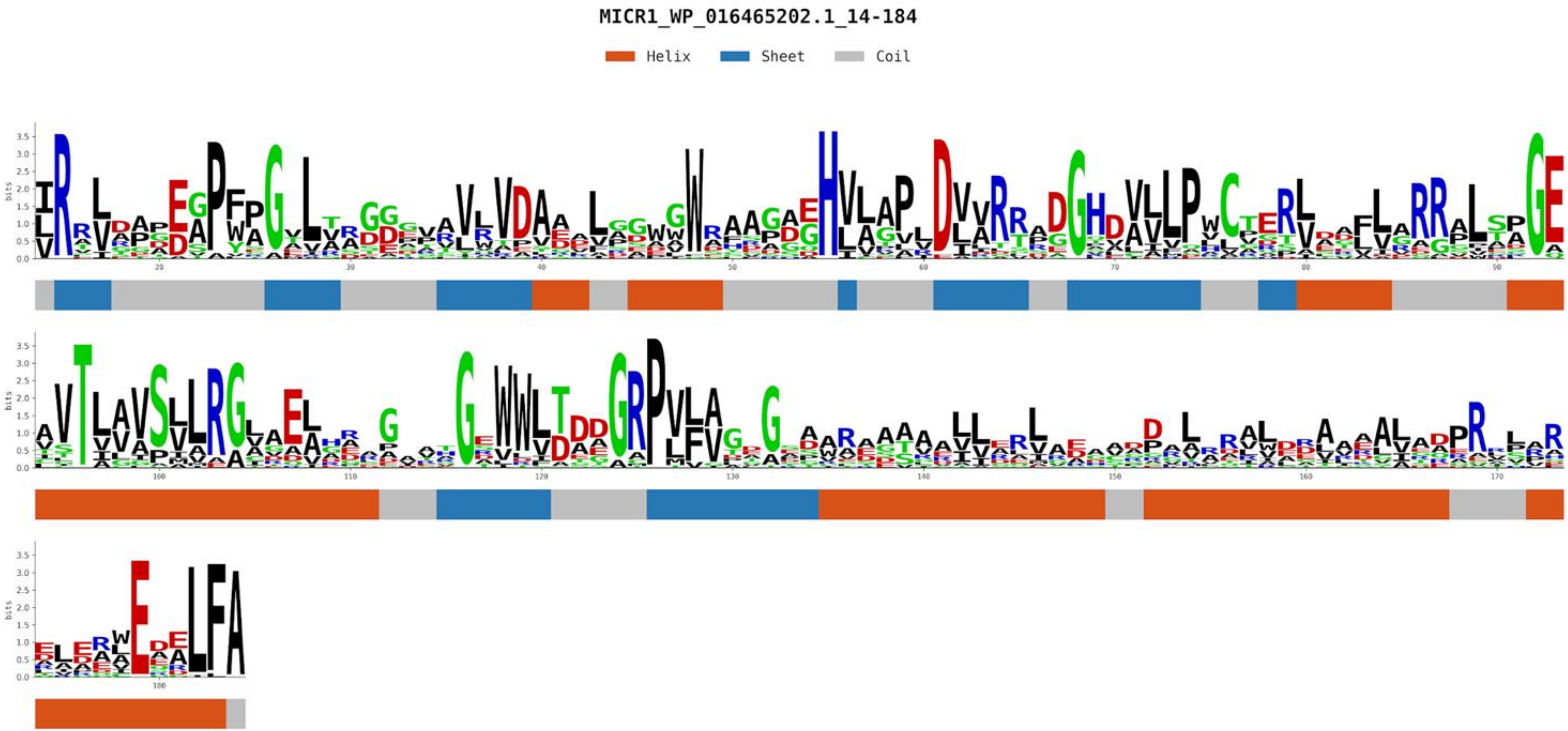


Only PKL fold recognized

POGI1_WP_013815849.1_1-438

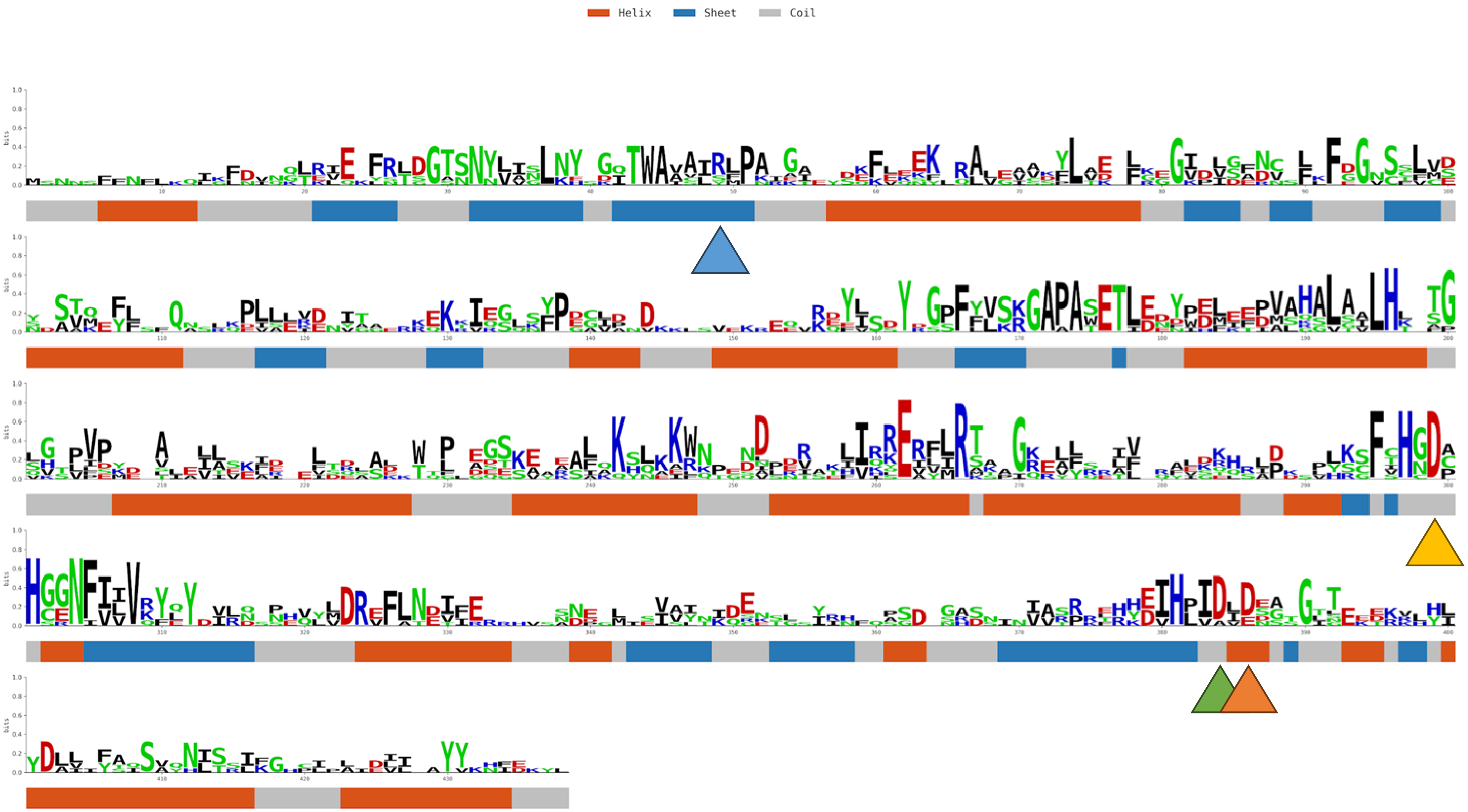

SEAE1_WP_147203046.1_59-310

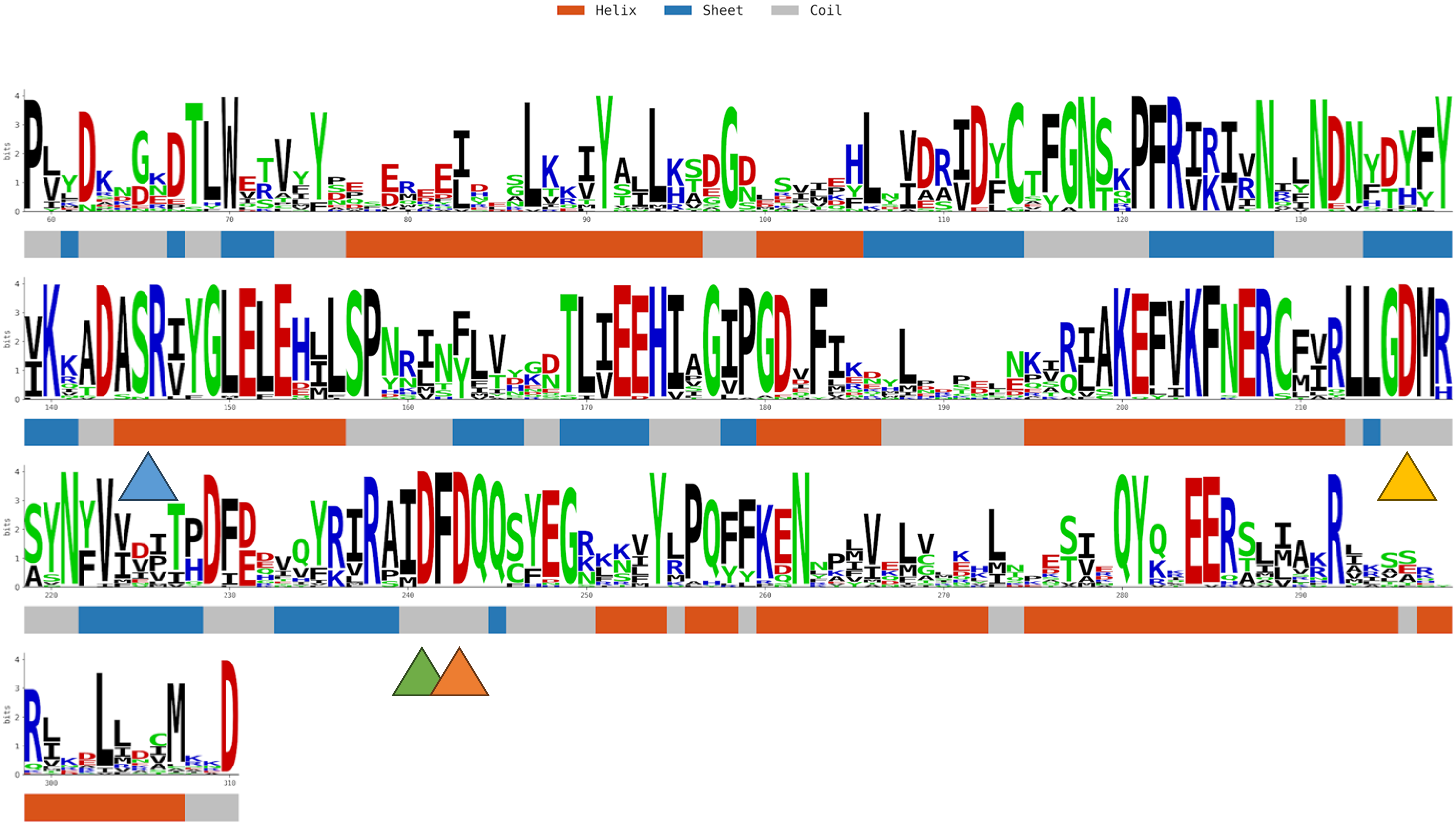

SELE1_WP_009753732.1_306-551

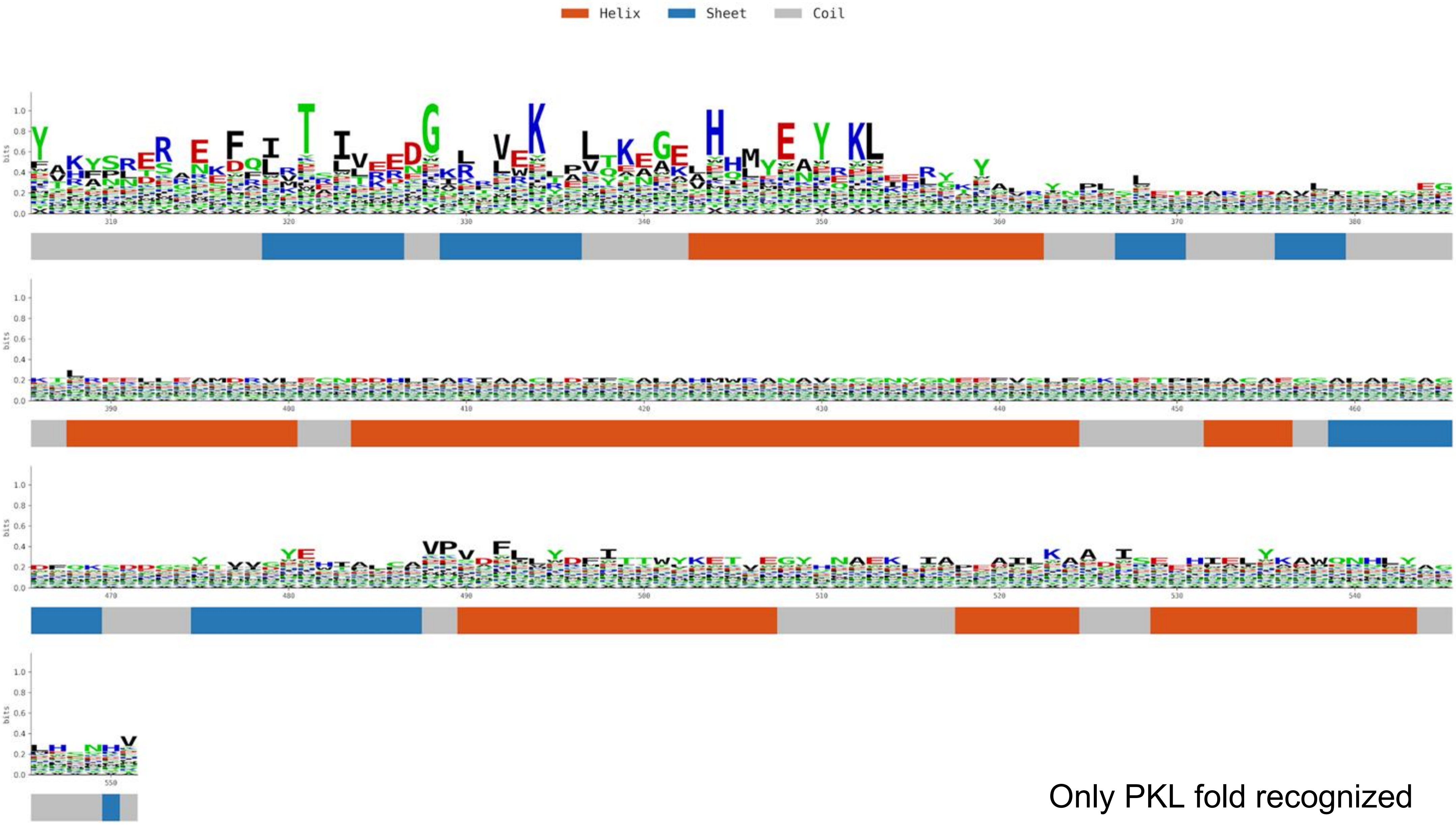


Only PKL fold recognized

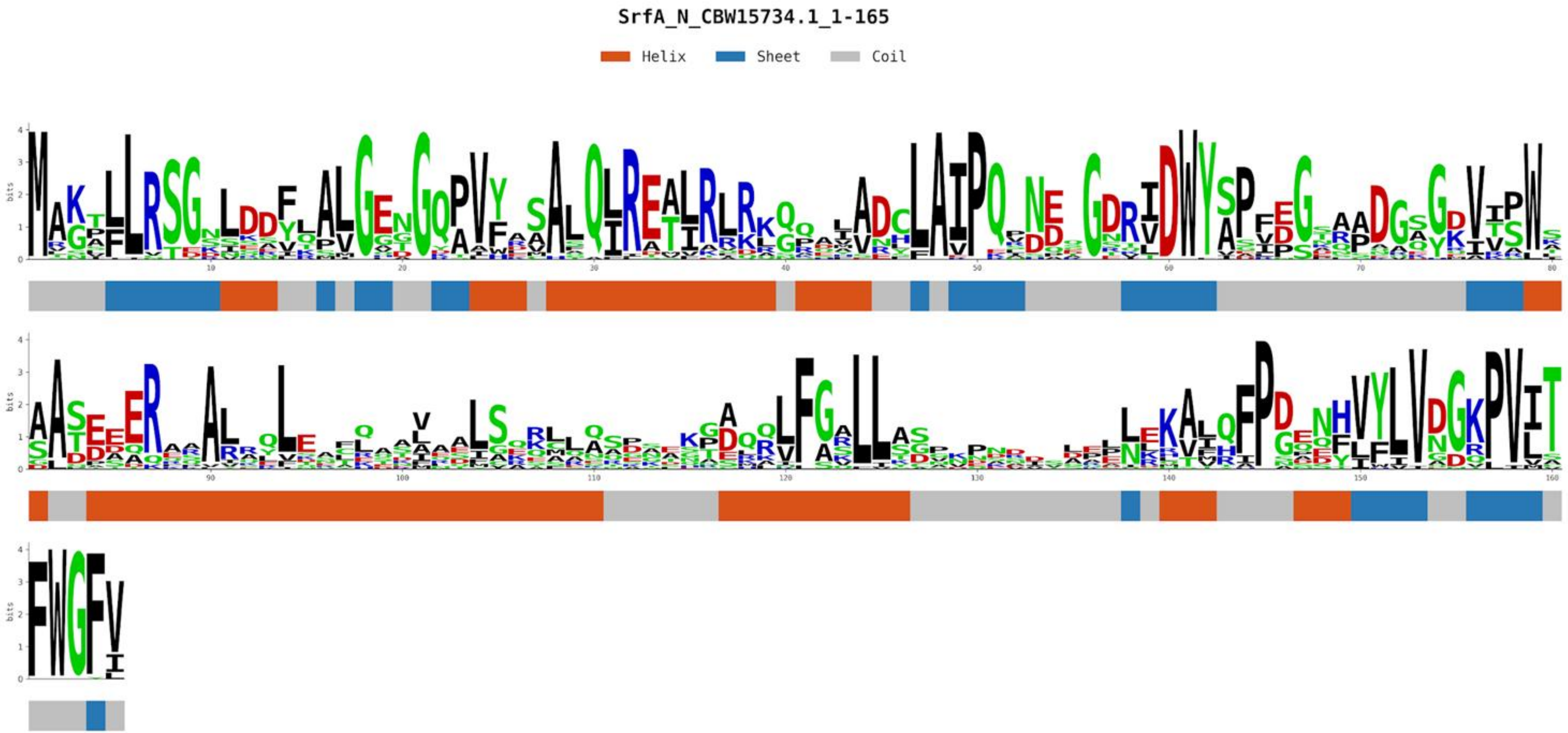


Only PKL fold recognized

STLA1_WP_200771915.1_22-337

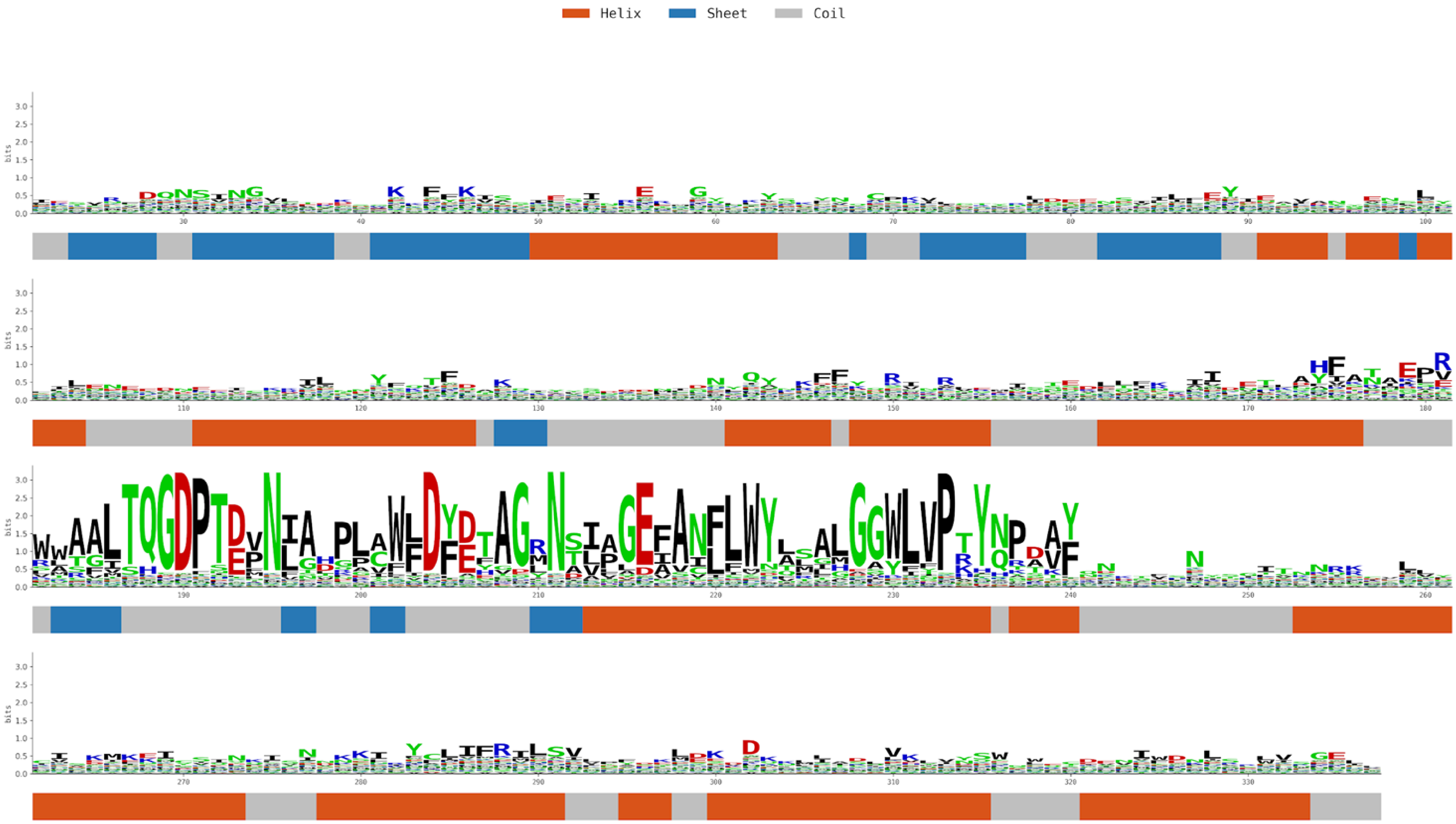

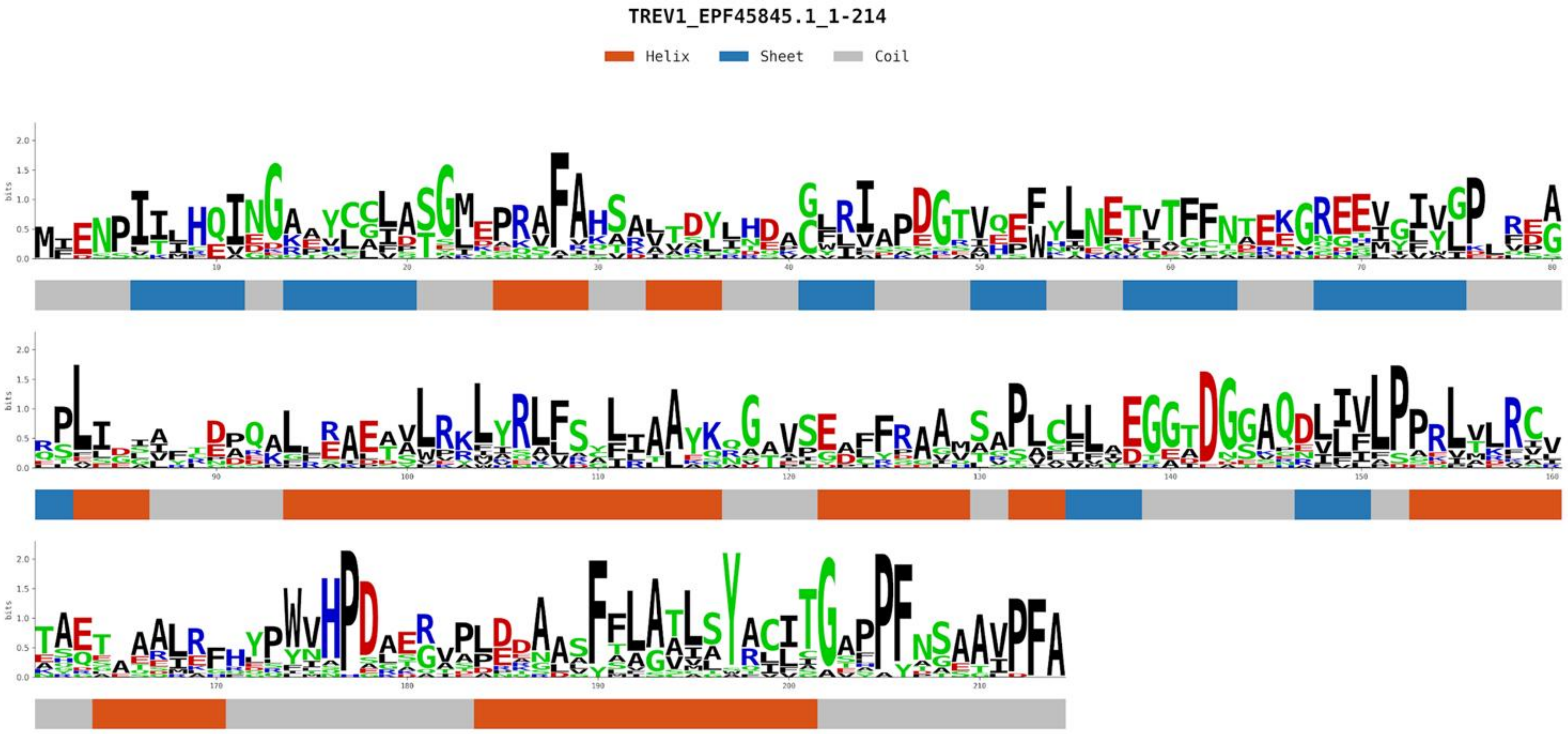


Only PKL fold recognized

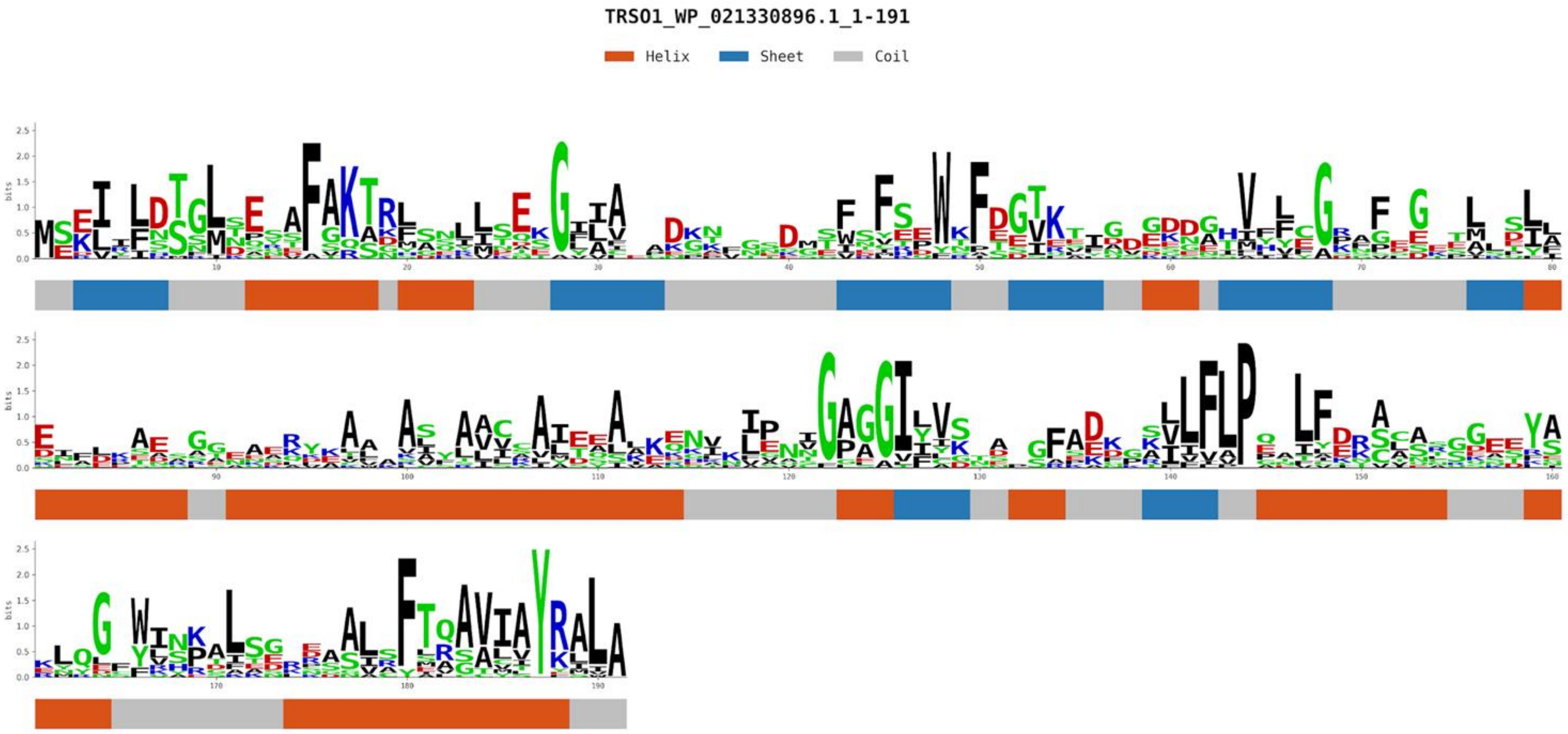


Only PKL fold recognized
